# Mutant p53 regulates cancer cell invasion in complex three-dimensional environments through mevalonate pathway-dependent Rho/ROCK signaling

**DOI:** 10.1101/2024.10.13.618100

**Authors:** Asja Guzman, Tatsuya Kawase, Alexander J. Devanny, Gizem Efe, Raúl Navaridas, Karen Yu, Kausik Regunath, Iris G. Mercer, Rachel C. Avard, Rafaela Muniz de Queiroz, Anil K. Rustgi, Laura J. Kaufman, Carol Prives

## Abstract

Certain mutations can confer neomorphic gain of function (GOF) activities to the p53 protein that affect cancer progression. Yet the concept of mutant p53 GOF has been challenged. Here, using various strategies to alter the status of mutant versions of p53 in different cell lines, we demonstrate that mutant p53 stimulates cancer cell invasion in three-dimensional environments. Mechanistically, mutant p53 enhances RhoA/ROCK-dependent cell contractility and cell-mediated extracellular matrix (ECM) re-organization via increasing mevalonate pathway-dependent RhoA localization to the membrane. In line with this, RhoA-dependent pro-invasive activity is also mediated by IDI-1, a mevalonate pathway product. Further, the invasion-enhancing effect of mutant p53 is dictated by the biomechanical properties of the surrounding ECM, thereby adding a cell-independent layer of regulation to mutant p53 GOF activity that is mediated by dynamic reciprocal cell-ECM interactions. Together our findings link mutant p53 metabolic GOF activity with an invasive cellular phenotype in physiologically relevant and context-dependent settings.

**Significance:** This study addresses the contribution of mutant p53 to the process of cancer cell dissemination in physiologically relevant three-dimensional environments – a key characteristic of metastatic disease. Several mutant p53 proteins display pro-oncogenic activity with respect to cancer cell invasion in 3D environments via mevalonate pathway-dependent Rho/ROCK signaling axis.

## Introduction

Despite significant progress in understanding the processes underlying primary tumor formation, our mechanistic understanding of metastatic disease, the aspect of cancer responsible for over 90% of cancer-related deaths ^1^, remains incomplete. Metastasis is a multi-step process, spanning a variety of events from cancer cell invasion at the primary tumor site to intra-and extravasation and outgrowth of metastatic colonies at distant organs ^2^.

The *TP53* gene is the most frequently mutated gene in human cancers with the majority of mutations being of the missense variety ^3^. Among these are a handful (∼6) of “hotspot” mutations that occur with extraordinarily high frequency. Beside diminishing the wild type activity of p53 (loss of function), there is growing evidence that certain mutant p53 proteins acquire neomorphic ‘oncogenic’ properties and promote the metastasis cascade through multiple avenues. Studies in mice showed that mutant p53 can contribute to tumor invasive behavior beyond the effects caused by wild-type or loss of p53 indicating pro-metastatic gain-of-function (GOF) activities ^4–11^. Mutant p53 was reported to promote cancer cell invasion through multiple mechanisms, including regulating epithelial-to-mesenchymal transition (EMT), regulating cell-extracellular matrix (ECM) interactions, promoting receptor tyrosine kinase (RTK) signaling, and various metabolic pathways ^12^.

While studies in mouse models have confirmed the physiological relevance of the pro-metastatic activities of mutant p53, the time scale needed for experiments in animals makes it challenging to decouple the contributions of cell proliferation and cell invasion. On the other hand, while there is a wealth of elegant studies demonstrating the effects of mutant p53 on cell locomotion in 2D and “pseudo-3D” settings, the mechanisms regulating cell migration in 2D and 3D scenarios are known to differ significantly ^13^. Which mutant p53-regulated pathways underlie the enhanced cancer cell locomotion in complex, physiologically relevant three-dimensional environments still remain to be further elucidated.

Cell locomotion in three-dimensional environments is dependent on establishment of reciprocal cell-ECM interactions and on the dynamic formation of cell protrusions that can be subdivided into actin filament-based structures (lamellipodia, filopodia and invadopodia), and the cell contractility-driven lobopodia and membrane blebs ^14^. These processes all involve the Rho family small GTPases, prominently Rho, Rac1 and Cdc42 ^15^. Cancer cells can dynamically switch between Rho-dependent bleb-mediated round cell invasion and the Rac1/Cdc42-dependent elongated mode that relies on filopodia and lamellipodia ^16^. The balance between Rho and Rac1/Cdc42 signaling is dictated by cell intrinsic properties as well as by the biomechanics of the surrounding ECM ^17,18^. The small GTPases are all subjects of tight spatiotemporal regulation by multiple signaling mechanisms ^19^. Beside regulation by the opposing actions of guanine nucleotide exchange factors (GEFs) and GTPase-activating proteins (GAPs) ^20^, the function of various small GTPases is also tightly regulated by isoprenylation, a posttranslational modification product of the mevalonate (MVA) pathway that is required for membrane tethering, effector binding and activity of the small GTPases ^21^.

Mutant versions of p53 have been previously linked to cell migration and RhoA-mediated signaling. Two different p53 hotspot mutant proteins were reported to support cancer cell migration in H1299 lung carcinoma cells by upregulating the recycling of the fibronectin-binding α5β1 integrin to the membrane through enhancing its interaction with a regulator of endocytic trafficking ^22^. Moreover, in breast cancer cells, the filopodia-inducing motor protein myosin-X was shown to be required for mutant p53-driven cell invasion, and to support cancer cell dissemination through transporting integrin receptors to actin-driven cell protrusions ^23^. Further, in mutant p53-driven mouse pancreatic ductal adenocarcinoma, activity of RhoA, a member of the Rho family of small GTP-binding proteins that controls actomyosin-driven cell contractility, was lost upon p53 knockout, indicating a role of mutant p53 in spatiotemporal regulation of RhoA activity ^24^. Mutant p53 was also demonstrated to upregulate the activity of the master regulator of the MVA pathway, the SREBP2 transcription factor by our group ^25^ and was subsequently reported to upregulate Yap/Taz activity through increased Rho GTPase signaling ^26,27^. A better understanding of the molecular basis of cancer invasion-supporting mutant p53 activities may facilitate identification of new vulnerabilities and instruct the design of novel therapies aimed at inhibiting cancer dissemination.

## Results

### Mutant p53 proteins regulate invasion into complex three-dimensional environments in cancer cell lines

To elucidate a potential positive contribution of mutant p53 to the process of local dissemination of cancer cells in soft tissues, we performed a 10-day *ex vivo* breast cancer lung colonization assay using native three-dimensional (3D) lung extracellular matrix (Fig. S1a). The native matrix was isolated from murine lung using a decellularization technique that preserves the biochemical composition, integrity and biomechanical characteristics of the ECM ^28^. To address potential mutant p53-dependent differences in cellular colonization in the decellularized lung tissue, we used the MDA-MB-231 triple negative breast cancer cell line (referred to as MDA-231 in the figures) that harbors mutant p53 (R280K) along with two p53 knock-out (KO) clones derived from MDA-MD-231 cells that were generated by CRISPR/Cas9 technology (Fig. S1b). After performing this assay, the H&E stained lung tissue slices revealed a striking loss of colonization such that the cells with R280K mutation displayed 2.6-7.3-fold greater numbers of colonies than the p53 KO cells (Figs. 1a). Although we also noted a decrease in percentage of proliferative KI67^+^ cells upon p53 deletion (Suppl. Fig. S1c), it is very similar between the two KO cell lines (factor 2.6 for KO4 and 1.9 for KO12), while the invasive capacity between the two KO cell lines depicts stronger differences, suggesting that the differences in colonization capabilities are not due solely to proliferation.

**Figure 1.**
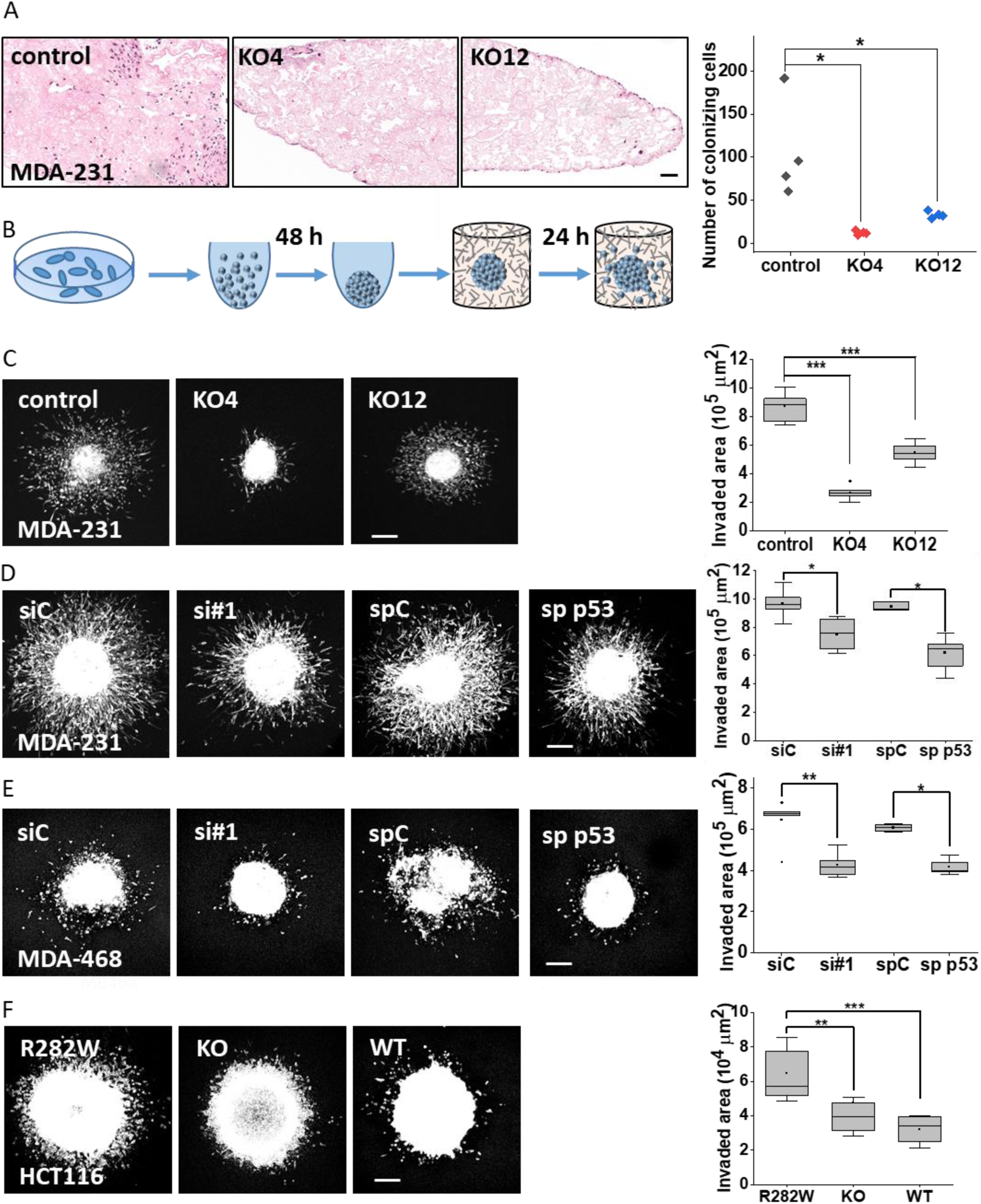
Mutant p53 supports cancer cell invasion in biomimetic 3D matrices. A) Representative H&E images (left) and quantification (right) of the number of colonizing MDA-MB-231 CRISPR control tumor cells harboring mutant p53 (R280K) or MDA-MB-231 CRISPR p53 KO cells *ex vivo* in decellularized lung tissue after 10 days, n = 4. Statistical significance was determined using Wilcoxon Rank Sum test, *p < 0.05. Scale bar = 50 µm. B) Experimental setup for a multicellular tumor spheroid (MTS) invasion assay involves spheroid formation under ultra-low adhesion conditions, embedding the fully formed MTS into a collagen I biomatrix and monitoring the extent of the invasion at 24 hours by confocal fluorescence imaging of the fluorescently labeled cells followed by quantification of the invaded area per MTS. (C-F) MTS invasion in collagen under mutant p53 depletion. Spheroids were prepared and subjected to the spheroid invasion assay in 1 mg/ml collagen gels. In each case at left are representative confocal fluorescence maximum projections of spheroids at t = 24 h (scale bar = 200 μm) and at right are the corresponding quantifications of MTS invasion. Invasion data in this and all following figures are presented as boxplots showing the median (black line) and second and third quartiles, with whiskers representing data from 5 % to 95 %. Squares indicate mean values. Dots show outliers. n signifies the number of spheroids per experimental condition unless stated otherwise. Statistical significance of these and all following invasion assays was determined using Wilcoxon Rank Sum test. (C) MDA-MB-231 CRISPR control, KO4 and KO12 cell lines. n = 11, 11, 12 for control, KO4 and KO12, respectively. (D) MDA-MB-231 parental cells transfected with non-targeting siRNA (siC), p53-targeting siRNA #605 (si#1), non-targeting smartpool control (spC) or p53-targeting smartpool siRNA (sp p53). n = 9, 4, 3, 6 for siC, si#1, spC and sp p53, respectively. (E) MDA-MB-468 cells under transient depletion of mutant p53 as in (D); n = 6, 6, 4 and 5 for siC, si#1, spC and sp p53, respectively. (F) HCT116 cell lines stably expressing the mutant p53 (R282W), no p53 (KO) or p53 wild-type (WT); n = 9, 10 and 10 for R282W, KO and WT cell lines, respectively.

To address whether the observed differences in lung tissue colonization were based mainly on a locomotive defect or were the result of a combination of locomotive and proliferative effects, we measured the proliferation of MDA-MB-231 CRISPR cells in 3D by performing a bioluminescent cell viability assay on multicellular tumor spheroids (MTSs) grown under ultra-low adhesion conditions for seven days (Suppl. Fig. S1d). This analysis showed that both MDA-MB-231 CRISPR p53 KO clones underwent significantly decreased proliferation in this experimental setting. The differences in proliferation between mutant p53-bearing vs. p53-depleted cells were smaller than the observed differences in lung tissue colonization efficiency (53 -34 % differences in average 3D MTS proliferation vs. 88 -69 % differences in average colonization). Taken together, these results indicate that the decreased number of p53 KO cells invading lung tissues after ten days of *ex vivo* culture reflects a locomotive defect caused by depletion of mutant p53 which is potentially exacerbated by differences in proliferative behavior.

To further decouple locomotive and proliferative effects and dissect mutant p53-dependent regulation of cancer cell motility in greater detail, we went on to perform short term MTS invasion assays in 3D collagen I hydrogels (Fig. 1b). This experimental system is biomechanically tunable, supports substantial multicellular invasion in a physiologically relevant yet well-controlled environment, and allows pharmacological treatments during the process of invasion. The MTS assay also allows assessment of invasion on a much shorter time scale (24 hours), thus minimizing the contribution of cell proliferation to the observed phenotypes. To this end spheroids were formed from various breast and colon cancer cell lines under transient and stable depletion of mutant p53 using a rapid (48 hours) spheroid formation protocol ^29^ and then fully embedded in low density collagen I matrices (1 mg/ml) for invasion. Samples were fixed 24 hours after embedding the spheroids and the invaded area per MTS was assessed from confocal fluorescence microscopy (CFM) images of the fluorescently labeled cells. To obtain cell proliferation data from each invasion assay, we used MTS size prior to onset of invasion as a proxy, since it fully mirrors the differences detected by the bioluminescent proliferation assay (compare Fig. S1e and f). As expected, proliferative differences between the CRISPR control and p53 KO cell lines are less pronounced after 48 hours than after the seven-day 3D culture protocol (compare Fig. S1d and g) and fluctuated between 13 and 46 % depending on the p53 KO cell line. Analysis of MTS invasion of these cell lines revealed that stable depletion of mutant p53 reduced spheroid invasion into collagen I matrices by a factor of two to four (Fig. 1c; Fig. Sh, i). This and all following invasion assays have been performed with 3-12 spheroids per condition with n signifying the number of spheroids used in the depicted experiment. Additional independent biological replicates are always shown in the Supplementary Materials section. The reduction of invasion far exceeded the extent of the detected proliferative differences. As further evidence for the impact of mutant p53 on invasion in this assay, transient siRNA-mediated depletion of mutant p53 using both a single targeting siRNA (referred to as “si#1”) or a mix of four targeting siRNAs (referred to as “smartpool” or “sp p53” throughout the manuscript) markedly reduced the invaded area of MDA-MB-231 spheroids (Fig. 1d; Fig. S1l, m; Western blot analysis of knockdown efficiency shown in Fig. S1j). While the MDA-MB-231 CRISPR cell lines showed significant proliferative differences in 3D MTS culture at various time points (Fig. S1d-g), transient knockdown of p53 in the MDA-MB-231 parental cell line did not cause significant differences in proliferation at t = 48 h (Fig. S1k). This indicates that the observed p53-dependent invasive phenotype is not primarily caused by proliferative effects. To exclude the possibility that the invasion-reducing effects caused by transient knockdown are the result of siRNA off-target effects, the p53-targeting and non-targeting siRNAs were transfected into the stable MDA-MB-231 control and KO cell lines and spheroid invasion was evaluated. While p53 knockdown expectedly reduced the spheroid invasion of the control cell line, it had no or only minor effects on invasive capacity in the KO cell line (Fig. S1n-p). This indicates that there are no major off-target effects with relation to invasion from the p53-targeting siRNAs that are used throughout this study.

To expand this study to other p53 missense mutants, we used MDA-MB-468 cells (referred to as MDA-468 in figures), another triple negative metastatic breast cancer cell line carrying the R273H hotspot p53 variant, and again saw that MTS invasion was markedly reduced upon transient siRNA-mediated depletion of mutant p53 without significant p53-dependent proliferative differences detected within this time scale (Fig. 1d; S2a-d).

As yet another approach, we utilized the HCT116 colon cancer cell line that harbors wild-type p53 and two isogenic versions that either express no p53 (KO cells) generated via CRISPR or the hotspot mutant p53 R282W generated using CRISPR/base editing. This provided an allelic series that both added another p53 mutation and expanded the study beyond breast cancer cells. We found that R282W-expressing cells displayed a significant invasive advantage over the p53 wild-type or p53-depleted cells in the MTS invasion assay while not displaying any proliferative differences on this time scale (Fig. 1f; Fig. S2e-h), indicating a true pro-invasive gain-of-function (GOF) role for mutant p53. To ensure that the observed GOF phenotype was not an artifact of the HCT116 R282W cell line, we transiently depleted mutant p53 from these cells using siRNA-mediated knockdown (Fig. S2i). Confirming that the increased invasion was due to the p53 knock-in R282W mutation, there was a marked reduction in invaded area of these cells; in contrast, no effect on the invasion of the p53-depleted HCT116 KO cell line transfected with the same siRNAs was evident (Fig. S2j-m). Taken together these results strongly support the hypothesis that the three p53 mutant proteins investigated here (R280K, R273H and R282W) exert an invasion-enhancing gain-of-function activity during the process of cancer cell dissemination in 3D collagen I environments that is independent of proliferative effects.

### Mutant p53 modulates cell morphology of 3D invading cancer cells

To elucidate the cellular underpinnings of the observed differences in invasion efficiency between mutant p53-carrying and -deficient cells, we analyzed cell morphology in 3D invading cancer cells. To this end we used the CRISPR MDA-MB-231 cell series since these cells displayed the widest distribution in cell morphology among the three cell lines used and so were the best suited to detects shifts in cell shape and cell protrusions. The MDA-MB-231 CRISPR MTSs were allowed to invade in collagen matrix for 48 hours (the longer time was needed to ameliorate cell crowding at the spheroid periphery), fixed, stained for actin cytoskeleton and subjected to imaging using fluorescence confocal microscopy. Images were taken at various positions of the invading spheroid, both at the very edge of the invading front as well as closer to the spheroid core. The majority of the cells analyzed here, however, was located at the invading front in order to avoid artifacts caused by cell crowding at positions closer to the spheroid core. Qualitative assessment revealed a shift towards an elongated cell morphology among the 3D invading p53-KO MDA-MB-231 cells in comparison to the mutant p53-carrying control cells (Fig. 2a, b). Quantitative analysis of cell morphology, as reflected by the cell circularity index, revealed a significant reduction of the median cell circularity in both p53 KO cell lines in comparison to the control cell line (Fig. 2c). The proportion of cells categorized with a circularity index ≥ 0.5 was reduced by approximately 50 % upon stable depletion of mutant p53.

**Figure 2.**
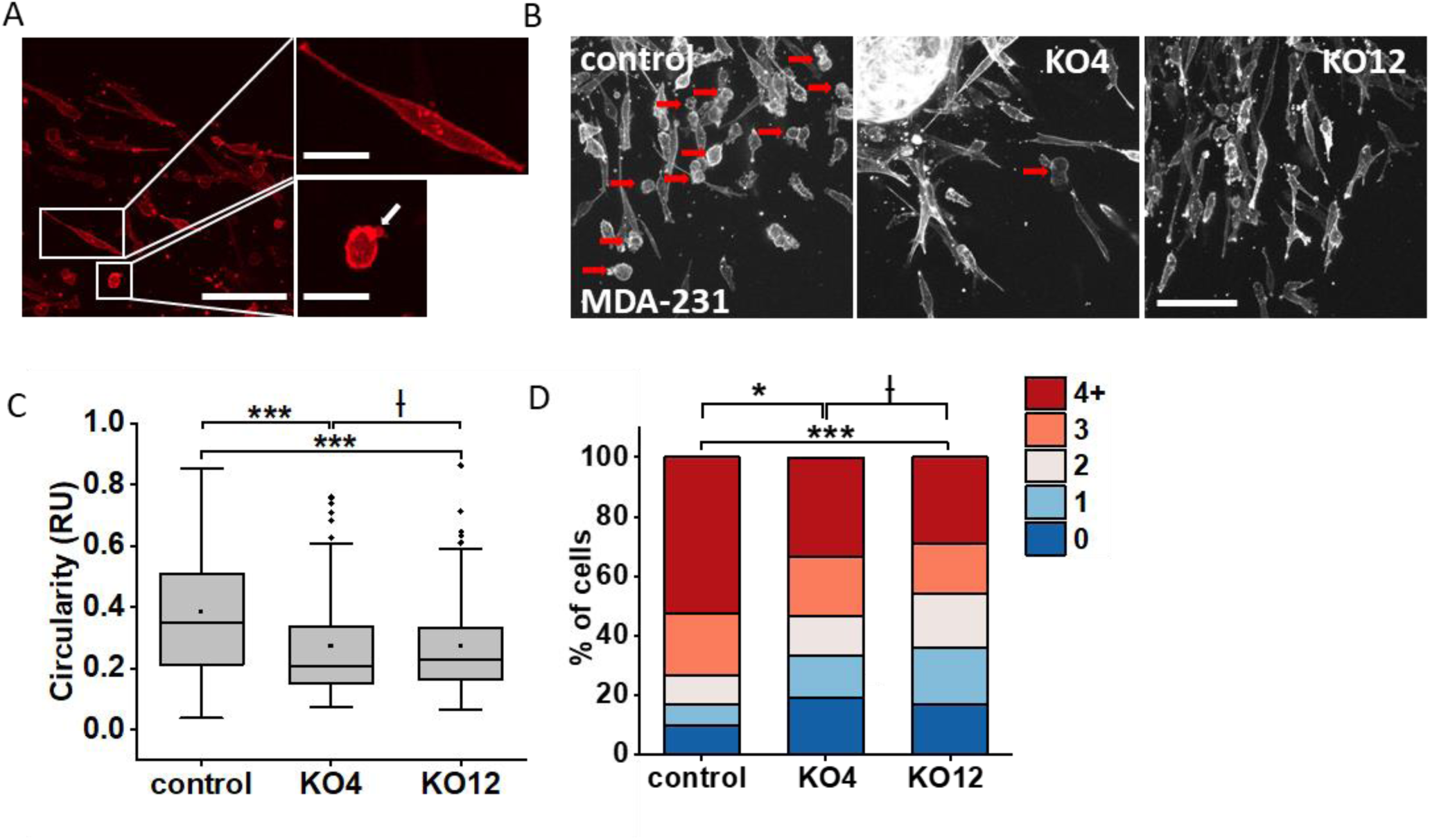
Mutant p53 regulates cell morphology of 3D invading cancer cells. A) Exemplary images of phalloidin-labelled filopodia/lamellipodia and bleb-bearing MDA-MB-231 CRISPR control cells after 48 h of MTS invasion in collagen I. White arrow indicate a membrane bleb. White squares indicate areas shown at a greater magnification in the adjacent images. Scale bar = 100 µm for the overview image and 50 µm for the high magnification insets. B) Representative 20x confocal fluorescence images of phalloidin-labelled MDA-MB-231 CRISPR control, KO4 and KO12 MTS after 48 h of invasion in collagen I. Red arrows indicate round cells. Scale bar = 50 μm. C) Quantification of cell circularity in invading cells of MDA-MB-231 CRISPR control, KO4 and KO12 MTS after 48 h of invasion in collagen I. The circularity indices are presented in a box plot. n = 100 cells per condition per experiment, data pooled from 2 individual experiments. D) Analysis of bleb-bearing cells in 3D collagen-invading MDA-MB-231 CRISPR control, KO4 and KO12 spheroids after 48 h of invasion in collagen I matrices. Bar plot shows percentage of cells bearing 0, 1, 2, 3 or ≥ 4 blebs per cell among the invading cell population. n = 100 cells per condition, data pooled from 2 individual experiments. Statistical significance in the distribution of number of blebs was established by Kolmogorov-Smirnov test.

Changes in cell circularity can indicate a shift in the balance between various signaling pathways that regulate cell contractility and cytoskeletal re-organization such as Rho/ROCK versus Rac signaling. The dynamic equilibrium between these pathways is crucial for establishment of different types of cell protrusions and for setting the preferred migratory mode in 3D environments. Accordingly, we asked whether mutant p53 is involved in setting the type and proportion of different cell protrusions established during 3D MTS invasion. The cell protrusions of invading cells (same data set as used for circularity analysis) were categorized as actin-polymerization-driven protrusions or membrane blebs (Fig. 2a), and the occurrence of such protrusions in the invading cells was determined. For membrane blebs, a cell protrusion governed by cell contractility and highly relevant for *in vivo* cancer cell invasion ^30–32^, the number of blebs per cell was determined, as our previous work suggested that the establishment of clusters of blebs is particularly relevant in 3D motility ^33^. This analysis revealed that the population of bleb-bearing cells was markedly reduced under stable depletion of mutant p53 in the MDA-MB-231 KO cell lines in comparison to the mutant p53-expressing MDA-MB-231 control cell line and the number of blebs per cell was lower (Fig. 2d). Taken together, these results indicate that mutant p53 regulates processes that support round cell morphology and the dynamic formation of migration-associated membrane blebs in 3D invading cancer cells, both of which are known to be regulated by the Rho/ROCK signaling cascade ^34^.

### Mutant p53 promotes 3D cancer cell invasion via the Rho/ROCK signaling axis

Since Rho/ROCK signaling is required not only for membrane bleb formation but also for cell contractility-dependent ECM re-organization ^35^, we performed a collagen gel contraction assay to compare overall levels of cell contractility between mutant p53-expressing and -depleted cells. This assay provides a readout reflecting the average 3D cellular contractility of a relatively large population of cells at 24 hours after embedding the cells in the matrix. MDA-MB-231 KO cells exhibited a significant reduction of average collagen gel contraction in comparison to the mutant p53-bearing control cells (Fig. 3a, b; Fig. S3a, b). Also MDA-MB-468 cells transiently depleted of mutant p53 showed a significant decrease in collagen gel contraction (Fig. 3c; Fig. S3c, d) that was also detectable at the initial stage of contraction (t = 4 h) thus indicating that the observed effect is independent of cell proliferation (Fig. S3e, f). Collagen contraction assays in the HCT116 cell lines, the least contractile among the three cell lines, also revealed a comparatively small, yet significant shift towards higher cell contractility upon introduction of the R282W mutant form of p53 that was above those observed for the KO or WT cells. (Fig. 3d; Fig. S3g, h).

**Figure 3.**
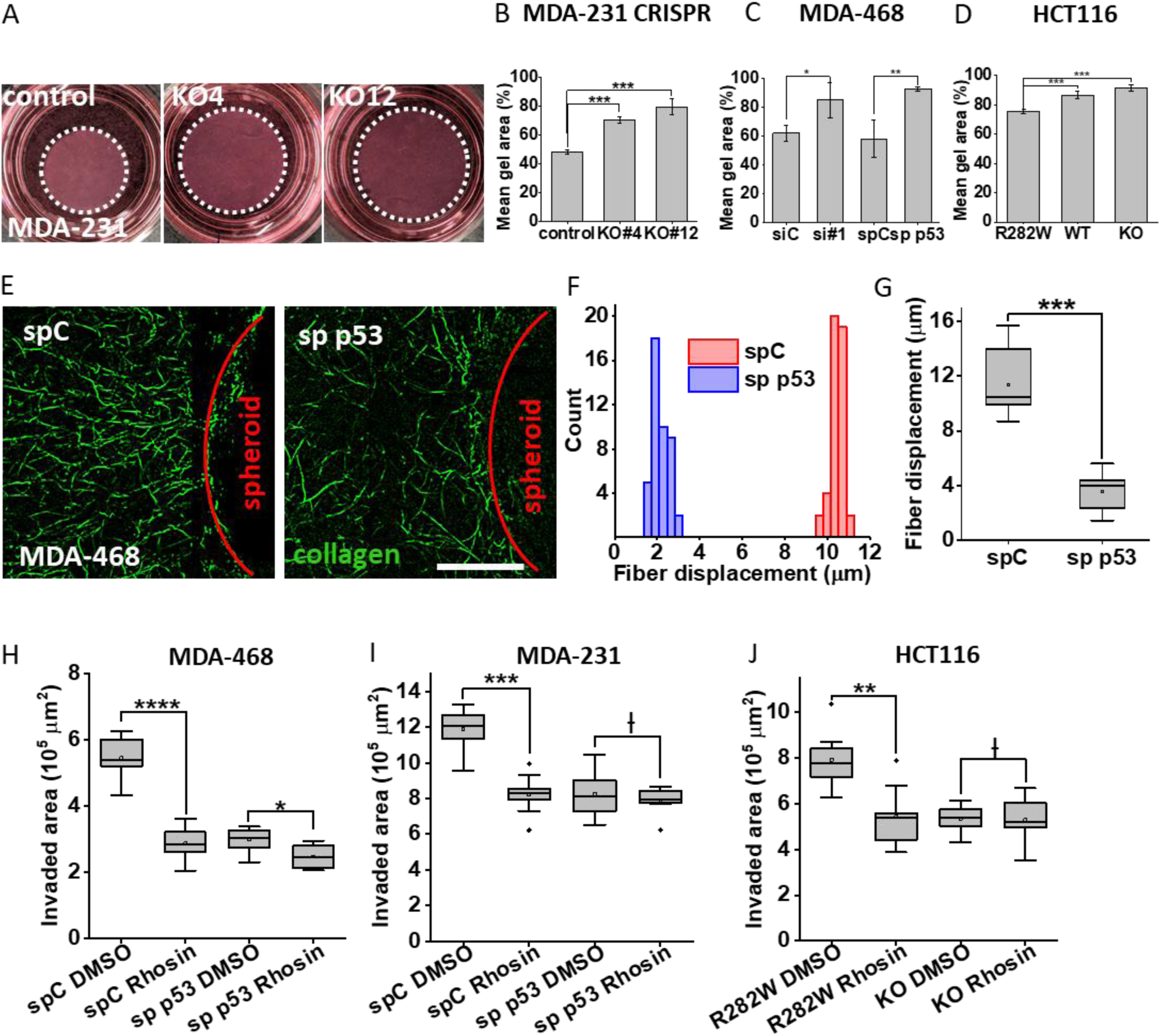
Depletion of mutant p53 interferes with cell-mediated ECM remodeling and collagen fiber alignment. A, B) Representative endpoint images (A) and quantification (B) from a collagen gel contraction assay with MDA-MB-231 CRISPR control, KO4 and KO12 cell lines. 1 mg/ml collagen I gels loaded with 1 x10^6^ cells/ml collagen were cast and allowed to contract unanchored immediately after polymerization. Gel area was measured before and after 24 h of contraction. Images show the 24 h time point. Histogram in (B) depicts average % gel area at the end point relative to initial gel area ± SD for these cell lines. n (number of collagen matrices per condition) = 3. Statistical significance for this and following collagen contractions assays was determined using two-sample unequal variance t-test. C) Quantification of a representative collagen contraction assay with MDA-MB-468 cells transiently transfected with p53-targeting siRNAs or the respective controls. Experiment was performed in quadruplicate. D) Quantification of a representative collagen gel contraction assay of HCT116 KO, p53 WT and R282W cell lines. Experiment was performed in triplicate. E-G) Analysis of collagen displacement by individual MTS. MTSs were prepared from MDA-MB-468 cells transiently transfected with a non-targeting or p53-targeting siRNA pool, embedded into fluorescently labelled 1 mg/ml collagen I matrices loaded with fluorescent beads and allowed to contract collagen for 2 h at 37 °C. E) Exemplary confocal fluorescence images of cell-mediated radial collagen fiber alignment at the edge of individual MTS either expressing or depleted of p53. Collagen is labeled in green, the MTS is visible as a black area devoid of collagen fibers with the border visualized by a red line. Scale bar = 20 µm. F, G) Quantitative analysis of collagen fiber displacement in the vicinity of an individual MTS as assessed through displacement of collagen-embedded fluorescent beads. Bead displacements after 2 h at 37 °C were quantified for one experiment (F) and for 3 independent biological replicates (G). n (total number of beads per condition) = 115 and 119 for spC and sp p53, respectively. Statistical significance was determined via Kolmogorov-Smirnov test. H-J) MTS invasion under pharmacological inhibition of Rho/ROCK signaling. Invasion quantification of MTS invasion assay under p53 depletion +/- pharmacological RhoA inhibition (15 µm Rhosin) in 1 mg/ml collagen. MTSs were formed from (H) MDA-MB-468 and (I) MDA-MB-231 cells transiently transfected with p53-targeting or control siRNAs or (J) HCT116 R282W and KO stable cell lines. n (MDA-MB-468) = 7, 8, 10, 10, 9, 10, 9 and 7 for siC DMSO, siC Rhosin, si#1 DMSO, si#1 Rhosin, spC DMSO, spC Rhosin, sp p53 DMSO and sp p53 Rhosin, respectively. n (MDA-MB-231) = 8, 9, 9, and 8 for spC DMSO, spC Rhosin, sp p53 DMSO and sp p53 Rhosin, respectively. n (HCT116) = 8, 9, 9, 10 for R282W DMSO, R282W Rhosin, KO DMSO and KO Rhosin, respectively.

To dissect the cell contractility phenotype in greater detail, we performed quantitative microscopy-based analysis of collagen contraction by individual spheroids. Here, MTSs from MDA-MB-468 cells transiently transfected with either non-targeting or p53-targeting smartpool siRNA were embedded in collagen gels containing fluorescently labeled beads and imaged first at the time of collagen gelation (t_0_ = 20 min), and then at a later time point when cell-mediated collagen-reorganization was readily visible but no invasion into the matrix had yet occurred (t_end_ = 120 min). As can be seen in the exemplary image shown in Fig. 3e, MDA-MB-468 MTS transfected with non-targeting siRNA displayed a radial alignment and accumulation of collagen fibers at the spheroid border that was not observed in the p53-depleted sample. To analyze this observation in a more quantitative manner, we measured fluorescent bead displacement reflecting the collagen fiber displacement in the vicinity of each spheroid (Fig. S3 i, j). The MTS collagen contraction analysis demonstrated a > 50 % reduction in average collagen fiber displacement upon transient mutant p53 depletion over the first two hours following collagen embedding of the spheroid, indicating that mutant p53 is required for full dynamic ECM reorganization, which precedes the process of cancer cell invasion into the ECM (Fig. 3f, g). Defects in collagen contraction can be caused by alterations in expression of integrin receptors that mediate cell adhesion to collagen. We used FACS to analyze the cell surface expression of β1 integrin, the high affinity collagen I-binding integrin receptor subunit, in all three cell lines. Neither transient p53 depletion in MDA-MB-468 cells nor stable p53 depletion in HCT116 and MDA-231 cells led to a decrease in cell surface presentation of β1 integrin indicating that it is not a defect in ECM-binding that is causing the reduced collagen contraction (Fig. S3k-p). Taken together, we have demonstrated a GOF of the three analyzed p53 mutants with respect to RhoA-driven cell contractility in 3D environments that is not caused by differences in integrin β1 presentation and can be observed in cells of different solid tumor origins, thus is not cell type-dependent.

To test if the reduction in invasiveness upon p53 depletion is mediated by the compromised Rho/ROCK signaling pathway, we performed MTS invasion assays with and without mutant p53 under pharmacological inhibition of the Rho/ROCK axis using two small molecule inhibitors, Rhosin and Y27632. Rhosin specifically inhibits the RhoA subfamily of small GTPases ^36^, while Y27632 inhibits Rho-associated, coiled-coil containing protein kinases ROCK1 and ROCK2 ^37^. If our hypothesis is correct that the mechanism behind mutant p53 pro-invasive activity is upregulation of Rho/ROCK signaling, then pharmacological inhibition of that signaling axis should not have notable effects on the invasion efficiency when mutant p53 levels are low. Indeed, MDA-MB-468 MTS invasion assays demonstrated that the reduction of invasion caused by transient mutant p53 depletion was not, or was only minimally, further enhanced by application of either Rhosin or Y27632, which in the mutant p53 scenario led to a significant decrease of invaded area (Fig. 3h; Fig. S4a-d). Confirmatory results were obtained with MDA-MB-231 and HCT116 spheroid invasion assays which also showed loss of sensitivity to Rho inhibition upon loss of mutant p53 expression (Fig. 3 i, j; Fig. S4e-i). From this we conclude that mutant p53 supports cancer cell locomotion in 3D collagen I environments through positive regulation of Rho/ROCK-dependent cellular mechanisms.

The contribution of Rho-dependent motility to the overall invasion efficiency depends on various cell-intrinsic characteristics and can be dictated by cell-independent factors, such as the biomechanical properties of the ECM. Differences in stiffness, pore-size and architecture of the ECM are sufficient to shift the balance between Rho-vs Rac1/Cdc42-driven migratory mechanisms ^38^ and thereby change the susceptibility of the cells to invasion inhibition through small molecules targeting these signaling pathways. We analyzed the pro-invasive effect of mutant p53 in collagen invasion assays using matrices that pose different requirements for Rho-mediated locomotive processes. Our initial MTS invasion assays demonstrating an invasion-supporting role for mutant p53 were performed in 1 mg/ml pepsin-treated (PT) collagen, a matrix that is very permissive for invasion due to its low stiffness (as characterized by storage modulus, G’ = 14.2 ± 0.65 Pa) and non-restrictive pore-size for the cell types used in this study (ξ = 5.2 ± 0.4 µm) (Fig. 4a, b, referred to as Low). To address whether mutant p53 regulates cell invasion under biomechanically more challenging conditions, we chose either a higher density collagen matrix (3 mg/ml collagen I) with greater stiffness (G’ = 166 ± 36.2 Pa) and smaller pore-size (ξ = 3.3 ± 0.1 µm) yet a comparable architecture of fibers to that in 1 mg/ml PT collagen (Fig. 4a, b, referred to as High). Additionally, we used a composite collagen/basement membrane extract (BME) matrix (1 mg/ml collagen I and 3 mg/ml BME, referred to as Mixed) which is more comparable in stiffness to low density collagen (G’ = 39.5 Pa ± 27.9 Pa) but presents an additional biomechanical challenge due to the nanoporous structure of BME filling the spaces between the collagen fibers. We also used a collagen matrix (4 mg/ml collagen I gelled at 22 °C) that exhibits higher stiffness (G’ = 192 ± 7.4 Pa) with non-restrictive pore-size (ξ = 4.8 ± 0.4 µm) and highly bundled fibers that are resistant to re-organization by cells (Fig. 4a, b, referred to as Bundled) ^39,40^. Efficient cancer cell invasion in high and low density collagen matrices was reported to require Rho/ROCK signaling ^35,41^. In accordance with these studies, we found that MDA-MB-231 spheroid invasion in low and high density collagen I as well as mixed collagen/BME matrices was dependent on both Rho/ROCK and Rac/Cdc42 signaling as shown by a significant reduction of invasion upon application of the Y27632 inhibitor in this experimental setting (Fig. 4c-e; Fig. S5a-f). In contrast, our data showed that invasion into bundled collagen relies predominantly Rac/Cdc42-driven locomotive strategies, as indicated by the insensitivity of MDA-MB-231 invasion to the ROCK inhibitor Y27632 while remaining sensitive to Rac/Cdc42 inhibition (Fig. 4f; Fig. S5g, h). In accordance with the Rho and ROCK inhibition results, transient depletion of mutant p53 in MDA-MB-231 cells did not have a significant impact on invasion of MTSs in bundled collagen matrix while it significantly reduced invasion in dense collagen matrices (Fig. 4g, h). Thus, we show here that the invasion-promoting GOF effect of mutant p53 depends on the biomechanical properties of the three-dimensional extracellular environment.

**Figure 4.**
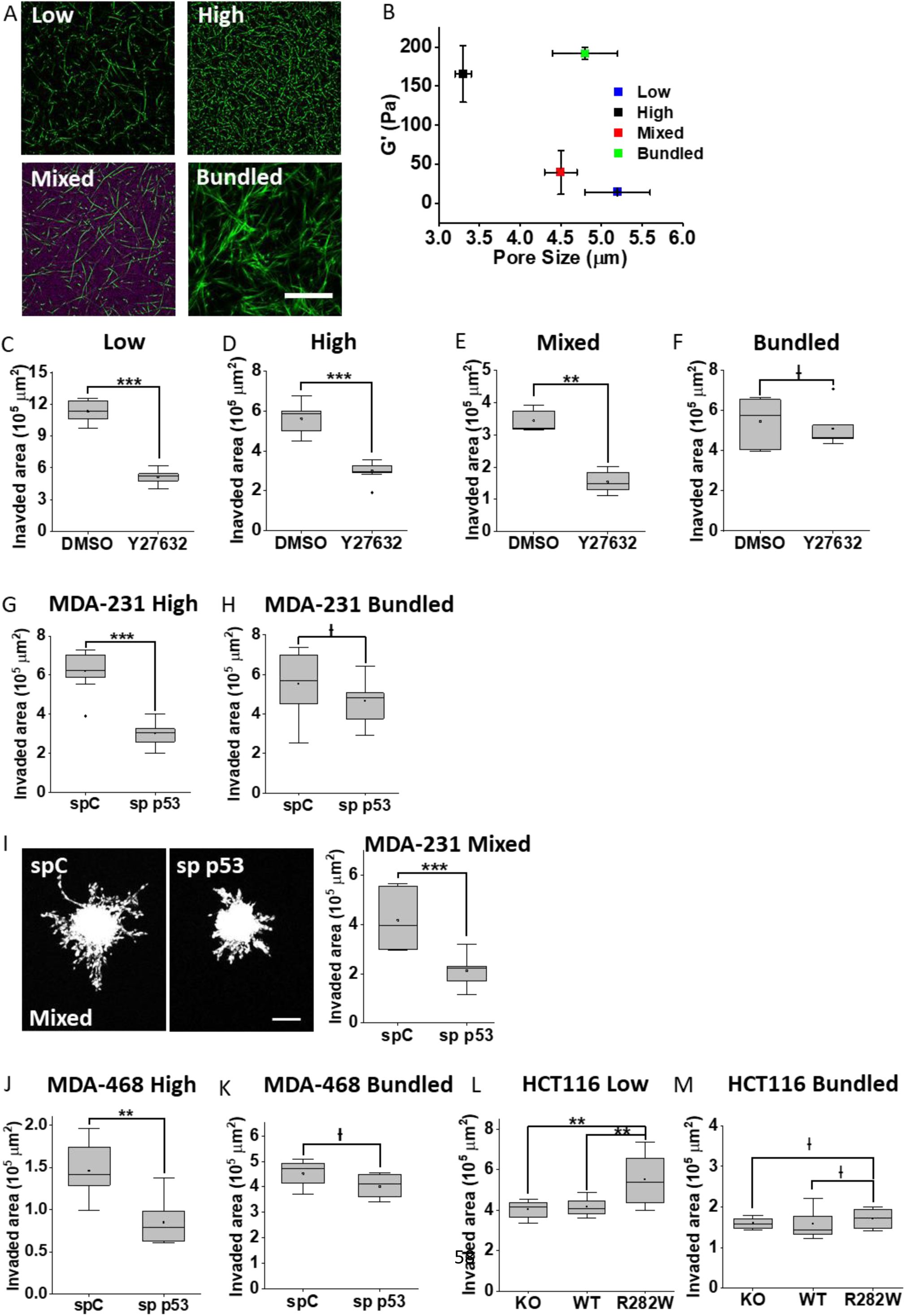
Pro-invasive GOF of mutant p53 depends on ECM biomechanics. A) Representative structured illumination microscopy (SIM) images of low concentration (1 mg/ml PT collagen, referred to as “low”) and high concentration (3 mg/ml PT collagen, referred to as “high”), representative confocal fluorescence Airyscan image of bundled collagen (4 mg/ml AS collagen gelled at 22 °C, referred to as “bundled”), and representative SIM image of mixed collagen/BME matrix (1 mg/ml PT collagen + 3 mg/ml BME, referred to as “mixed”; collagen is shown in green, BME in red). Scale bar = 50 μm. B) Biomechanical properties of the various matrices shown in panel A) are shown as a function of storage modulus G′ (Pa) ± SD and the average pore size ξ (μm) ± SD over n ≥ 5 gels. Pore size refers to pores between collagen fibers in all matrices including the composite collagen/BME matrix. Pore size of the BME component is not reflected in this analysis. Error bars are not visible if smaller than symbols. C-F) Representative MTS invasion assays of MDA-MB-231 cells in (C) low, (D) high, (E) mixed and (F) bundled collagen matrices under pharmacological inhibition of Rho/ROCK pathway. MTSs were formed from untreated cells and embedded into collagen gels supplemented with 10 µM Y27632 or DMSO, respectively, for the entire duration of the invasion assay. n (low) = 8 spheroids per condition; n (high) = 10 and 9 spheroids for DMSO and Y27632; n (mixed) = 5 and 8 spheroids for DMSO and Y27632; n (bundled) = 6 spheroids per condition. Experiments presented in panels C-F were performed using MTSs from the same biological replicate. G) Invasion quantification of a representative MDA-MB-231 spheroid invasion assay under transient p53 depletion in high concentration collagen matrix. n = 10 spheroids for spC and sp p53, respectively. H) Invasion quantification of a representative MDA-MB-231 spheroid invasion assay under transient p53 depletion in bundled collagen matrix. MTSs used in this invasion assay originate from the same biological replicate as in panel G. n = 8 and 11 spheroids for spC and sp p53.I) Representative MTS invasion assay in composite collagen/BME matrix under p53 depletion. Representative confocal fluorescence maximum projections of p53 expressing (spC) vs depleted (sp p53) MTSs invading in mixed collagen matrix at 24 h are shown on the left, the quantification of a representative invasion assay on the right. Scale bar = 200 μm. n = 7 and 9 for spC and sp p53, respectively. J, K) Invasion quantification of a representative MTS invasion assay with stable HCT116 KO, WT and R282W cell lines in J) low concentration and K) bundled collagen matrices. n (low) = 11, 8 and 12 for KO, WT and R282W, n (bundled) = 9, 11 and 8 for KO, WT and R282W, respectively.

We also addressed the question of whether mutant p53 facilitates only individual cell invasion in 3D environments or whether it also enhances collective cancer cell invasion. Collective invasion, a migratory mode of particular relevance for metastatic dissemination *in vivo* ^42^ relies strongly on Rho/ROCK-mediated actomyosin contractility ^43^. MDA-MB-231 cells adopt this invasive strategy when exposed to a composite collagen/BME hydrogel, referred to as “Mixed” ^44^, in which the cells are surrounded by a biomechanically challenging nanoporous laminin-rich matrix that also contains fibrillar collagen I, providing migration-supporting physical cues and biochemical ligands. Here, too, the presence of mutant p53 significantly enhanced collective MDA-MB-231 invasion (Fig. 4i; Fig. 5i-k) supporting the hypothesis that the invasion-supporting effects of mutant p53 are manifested in environments which require Rho/ROCK-dependent cellular mechanisms for efficient locomotion. Also in MDA-468 cells and in HCT116 isogenic colon carcinoma cell lines the invasive advantage contributed by presence of mutant p53 as observed in low and high density collagen was fully lost in the bundled collagen matrix (Fig. 4 j-m; Fig. S5l, m). This strongly supports our hypothesis that mutant p53 is crucial for Rho/ROCK-dependent individual and collective invasive behavior across cancer cells of different origin.

**Figure 5.**
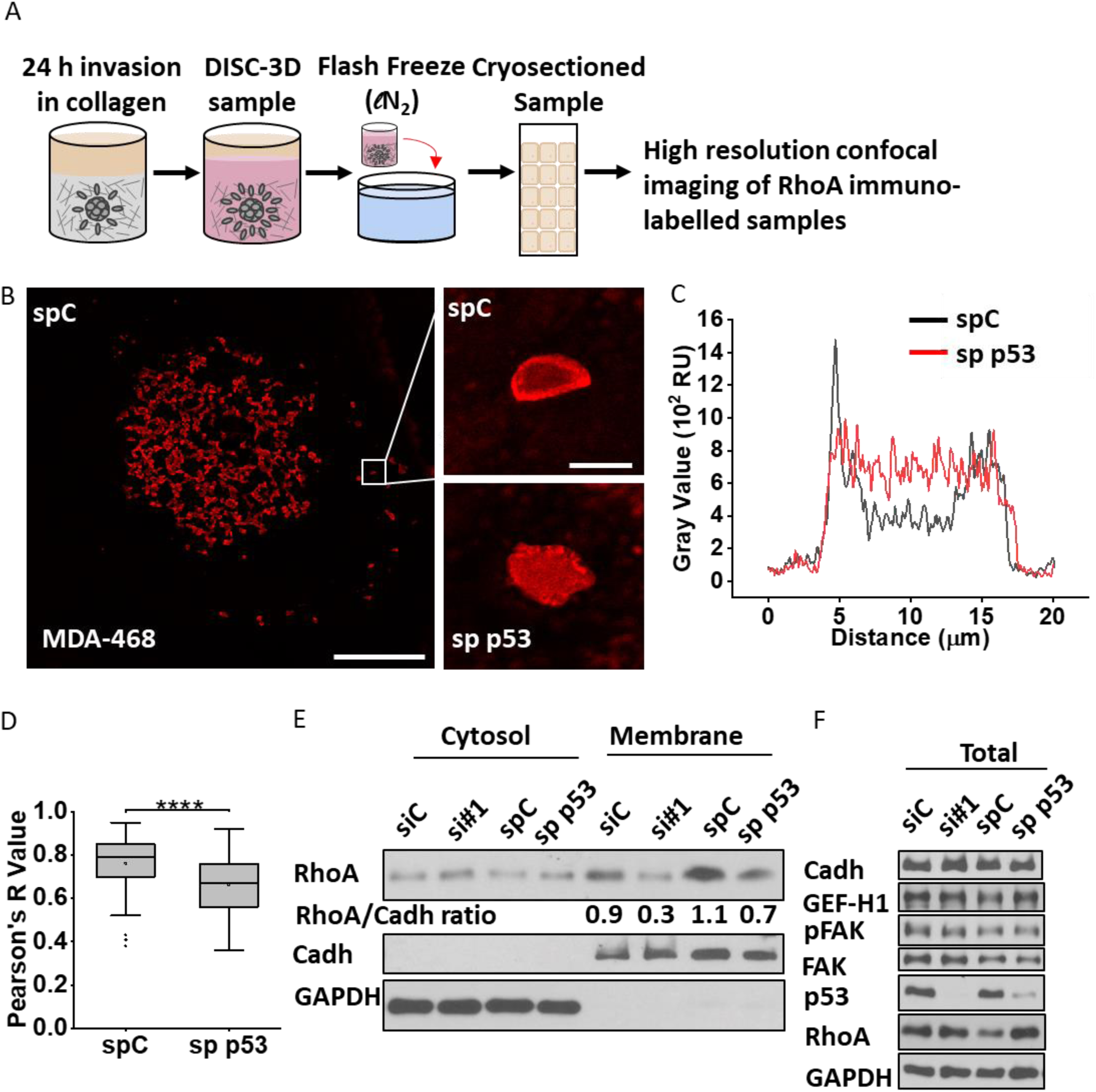
Mutant p53 regulates subcellular RhoA localization. A) Experimental setup for analysis of endogenous RhoA distribution in 3D invading spheroids (adapted from ^47^). Spheroids are prepared from MDA-468 cells and subjected to the MTS invasion assay in 1 mg/ml PT collagen. After 24 h samples are treated according to the DISC-3D protocol, cryosectioned, subjected to immunohistochemical fluorescent labeling of RhoA prior to high resolution confocal fluorescence imaging. B) Exemplary confocal fluorescence images of RhoA subcellular distribution in representative cells invading into collagen I gels from MDA-MB-468 spheroids transiently transfected with either p53-targeting siRNA pool (sp p53) or the respective non-targeting control (spC) obtained following the DISC-3D protocol described in A). Scale bar = 200 µm on the low magnification image showing the entire invading spheroid, and scale bar = 10 µm on the two insets at right showing individual cells. C) Exemplary RhoA fluorescence intensity line plot obtained from the two individual cells shown in high magnification in B). D) Quantification of subcellular RhoA distribution in invading MDA-MB-468 spC and sp p53 cells. Data is shown as Pearson’s correlation coefficient of RhoA with β1 integrin used as a cell membrane marker. n (number of quantified cells) = 100, data from 3 independent biological replicates. Comparison of Pearson’s R values for RhoA and β1 integrin in mutant p53 versus p53-depleted cells was carried out using Wilcoxon Rank-Sum test. E) Western blot analysis of subcellular RhoA localization as assessed by cell fractionation of MDA-MB-468 cells under transient siRNA-mediated depletion of mutant p53. 48 h post transfection cells were seeded for 90 min on collagen-coated plates, harvested and lysates were subjected to cell fractionation. A representative experiment is shown. Assessment of relative RhoA amounts in membrane fractions by densitometric analysis of RhoA signal intensity normalized to the cadherin signal intensity in the respective fraction (numbers shown below the RhoA panel). Total cell lysates shown in panel F). F) Western Blot analysis of Cadherin, GEF-H1, phospho-FAK and total FAK, as well as RhoA protein levels in total cell lysates of MDA-MB-468 cells under transient p53 depletion treated as in E.

### Mutant p53 upregulates the mevalonate pathway in order to produce isoprenoids for RhoA membrane attachment

Next we addressed the question of which aspect of Rho/ROCK signaling is regulated by mutant p53. Since RhoA localizes to the plasma membrane in order to bind its effectors and mediate its biological functions ^45,46^, we sought to analyze potential p53-dependent differences in RhoA localization in a 3D scenario during the process of cell invasion into collagen. For this we prepared cryosections from 3D collagen invading MDA-MB-468 MTSs under transient p53 depletion using our newly developed DISC-3D protocol ^47^, and subjected them to immunohistochemical labeling of endogenous RhoA and integrin β1 (as a membrane marker) (Fig. 5a). Imaging of individual cells from the invading front revealed differences in RhoA distribution as can be seen in the exemplary image and line plot of RhoA signal intensity across a representative cell (Fig. 5b, c). The line plot shows that while the mutant p53-expressing cells displayed peaks of RhoA intensity on the membrane with a drop of intensity in the cytosolic region, the p53-depleted cell displayed uniform RhoA distribution across the cell body without detectable accumulation of RhoA at the membrane (Fig. 5c). Co-localization analysis between RhoA and the integrin receptor as reflected in the Pearson’s correlation of signal intensity between these two proteins revealed a significant reduction in the coefficient indicating reduced RhoA localization to the plasma membrane in p53-depleted collagen-invading MDA-MB-468 cells (Fig. 5d). As another means of evaluating RhoA distribution in p53-depleted MDA-MB-468 cells, we performed cellular fractionation which confirmed that depletion of mutant p53 by either a single or a pool of p53-targeting siRNAs decreased localization of RhoA to the plasma membrane, while not decreasing overall RhoA levels (Fig. 5e, f; Fig. S6).

RhoA localization to the membrane is dependent on post-translational lipidation through a covalent transfer of a geranylgeranyl-moiety, a product of the MVA pathway that we and others have previously reported to be positively regulated by mutant p53 ^25,27^. To establish if there is a functional connection between the mevalonate pathway and cancer cell dissemination in our experimental setting, we performed MTS collagen invasion assays under pharmacological inhibition of the mevalonic acid pathway targeting the pathway at various points: Simvastatin is an inhibitor of the rate-limiting enzyme HMGCR ^48^, Fatostatin inhibits maturation and nuclear translocation of SREBPs ^49,50^ and GGTI-2133 blocks the enzyme geranylgeranyltransferase ^51^. All three inhibitors significantly reduced MDA-MB-468 MTS invasion to a degree comparable with p53 depletion (Fig. 6a). Simvastatin-induced reduction of MDA-MB-468 MTS invasion could be rescued by addition of exogenous geranylgeranylpyrophosphate (GGPP) indicating that this post-translational modification is required for the full extent of 3D invasion in these cells (Fig. 6b; Fig. S7a). Simvastatin also significantly inhibited invasion in MDA-MB-231 and HCT116 cell lines and this reduction could be rescued by exogenous GGPP (Fig. 6c-d; Fig. S7b, c), suggesting that protein lipidation is generally important for the process of 3D cancer cell dissemination across these different cell lines.

**Figure 6.**
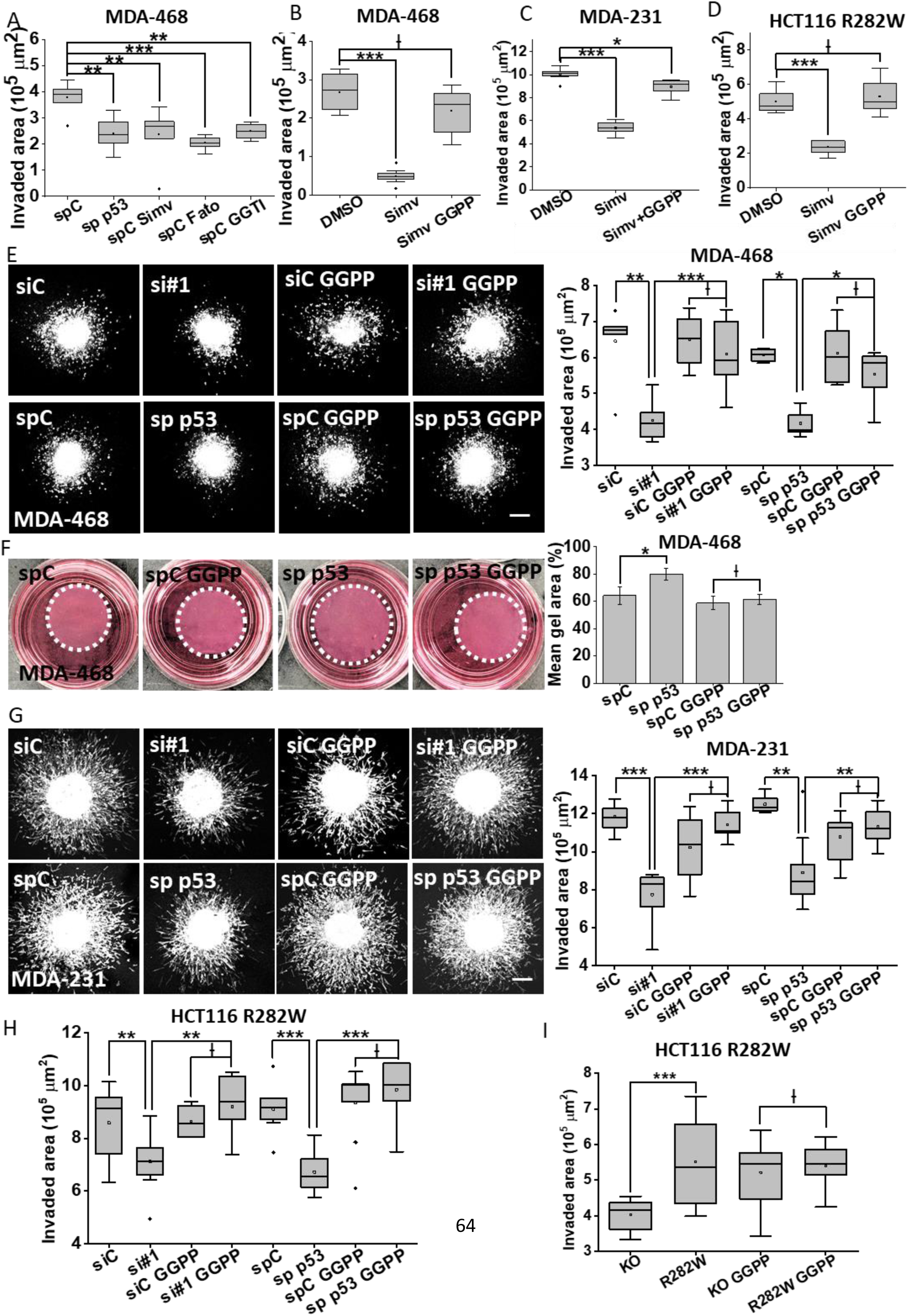
Mutant p53 promotion of 3D invasion is mediated by the mevalonate pathway. A) Comparative analysis of invaded area in MDA-MB-468 under transient depletion of mutant p53 or under pharmacological inhibition of mevalonic acid (MVA) pathway. n = 7, 8, 7, 6, 6 for spC DMSO, sp p53 DMSO, spC Simvastatin, spC Fatostatin and spC GGTI-2133, respectively. Statistical significance of this and other MTS invasion assays in this figure was determined using Wilcoxon Rank Sum test. B-D) MTS invasion of different cancer cell lines under pharmacological inhibition of MVA pathway (Simvastatin) +/- geranylgeranylpyrophosphate (GGPP) add-back. n (MDA-468) = 11, 9, 12 for DMSO, Simvastatin-and Simvastatin/GGPP-treated conditions. n (MDA-231) = 8, 11, 8 for DMSO, Simvastatin-and Simvastatin/GGPP-treated conditions, respectively. n (HCT116) = 6, 7, 6 for DMSO, Simvastatin-and Simvastatin/GGPP-treated conditions, respectively. E, G) Representative confocal fluorescence maximum projection images and MTS invasion analysis of MDA-MB-468 and MDA-MB-231 under mutant p53 depletion +/- GGPP add-back. MTSs prepared from cells transfected with either non-targeting (siC, spC) or p53-targeting (si#1, sp p53) siRNAs were subjected to invasion assay in collagen gels +/- GGPP. n (MDA-MB-468) = 6, 10, 6, 8, 4, 7, 5, 6 for siC untreated, sic GGPP, si#1 untreated, si#1 GGPP, spC untreated, spC GGPP, sp p53 untreated and sp p53 GGPP, respectively. n (MDA-MB-231) = 10, 8, 11, 10, 6, 11, 10, 10 for siC untreated, sic GGPP, si#1 untreated, si#1 GGPP, spC untreated, spC GGPP, sp p53 untreated and sp p53 GGPP, respectively. F) Representative images and quantification of a collagen contraction assay with MDA-MB-468 cells under transient p53 depletion +/- GGPP. MDA-MB-468 transiently transfected as described in E were embedded in 1 mg/ml collagen I gels +/- GGPP. Bar graph depicts average % gel area at the end point relative to initial gel area ± SD.Experiment was performed in quadruplicate. Statistical significance was determined using two-sample unequal variance t-test. H) MTS invasion analysis of HCT116 R282W under transient depletion of mutant p53 +/- GGPP. n = 8, 8, 9, 9, 10, 9, 8, 7 for siC untreated, sic GGPP, si#1 untreated, si#1 GGPP, spC untreated, spC GGPP, sp p53 untreated and sp p53 GGPP, respectively. I) MTS invasion analysis of stable HCT116 WT and KO cell lines +/- GGPP in comparison to HCT116 R282W cell line. n = 8, 10, 11, 10, 12 and 6 for WT DMSO, WT GGPP, KO DMSO, KO GGPP, R282W DMSO and R282W GGPP, respectively.

The next question was whether the invasion-enhancing activity of mutant p53 is mediated through providing non-sterol isoprenoids for post-translational modification of cell motility-related proteins. To answer this question, we supplemented MTSs formed from mutant p53-carrying and -depleted MDA-MB-468 cells with exogenous GGPP for the duration of the invasion assay. Strikingly, adding GGPP was sufficient to restore the 3D invasiveness of MDA-MB-468 cells under depletion of mutant p53 by either a single p53-targeting or a p53-smartpool siRNA (Fig. 6e; Fig. S7d, e). Additionally, the cell contractility of MDA-MB-468 cells under p53-depletion as assessed through a collagen contraction assay was restored upon supplementation with exogenous GGPP during collagen contraction assay (Fig. 6f; Fig. S7f). In full agreement with the result obtained with MDA-MB-468 cell line, MTS invasion of p53-depleted MDA-MB-231 and HCT116 cells was also rescued by adding back GGPP (Fig. 6g-I; Fig. S7g, h). Interestingly, supplementation with exogenous GGPP in HCT116 WT p53 cells increased the MTS invasion to the level observed for HCT116 R282W mutant p53 cells (Fig. S7h), providing further evidence for true GOF activity of mutant p53 mediated through regulation of post-translational lipidation. Taken together, these results demonstrate that the invasion-supporting GOF activity of three different p53 mutants is mediated through positive regulation of the MVA pathway and specifically through regulation of protein geranylgeranylation.

Analysis of gene expression of the MVA pathway enzymes in p53-depleted MDA-MB-468 spheroids revealed a significant reduction of expression of several genes in the pathway, which confirmed and extended our previously reported findings in a different cellular context (Fig. S8a) ^25^. Moreover, comparison of MVA gene expression in MTSs formed from the stable isogenic HCT116 cell lines revealed that expression of those genes is significantly higher in MTSs expressing the R282W mutant p53 than in those expressing the wild-type form or being depleted of p53 (Fig. S8b). Note that, consistent with our previous study showing that wild-type p53 suppresses MVA pathway gene expression ^52^, levels of MVA pathway gene mRNA were consistently lower in the wild-type p53 HCT116 cells than in the cells lacking p53 expression, while levels of the cholesterol transport protein ABCA1 were the lowest in mutant p53-expressing cells. This result supports the conclusion that that the ability of mutant p53 to regulate transcription of MVA pathway genes underlies the observed MVA-dependent regulation of the 3D cancer cell invasion and that it is not a cell type-dependent phenomenon.

### Mutant p53 mevalonate pathway gene target IDI-1 stimulates cell migration

We performed *in silico* analysis of MVA gene expression in breast cancer patients from the TCGA data base and found that several genes of the MVA pathway, among them IDI-1 and ACAT2, exhibit significantly higher expression in the tumor tissue in comparison to the normal tissue (Fig. 7a; Fig. S9a). Moreover, we observed that the molecular subtypes that correlated with higher rates of p53 mutations, such as the basal-like and Her2-enriched subtypes ^53^ have higher expression levels of IDI-1 (Fig. 7a). In fact, IDI-1 expression in breast cancer patients is higher in patients harboring missense p53 mutations when compared to those carrying wild type or no p53 (i.e. truncation mutations) with this trend being present across all tumor stages (Fig. 7b). A similar trend was observed for ACAT2 and also FDPS, another member of the MVA pathway involved in generation of geranyl pyrophosphate (Fig. S9b, c). Analysis of the overall breast cancer patient survival from the same cohort indicated an intermediate but highly significant decrease in survival in patients displaying high IDI-1 expression in tumor tissues, as shown in the Kaplan-Meier curve in Fig. S9d.

**Figure 7.**
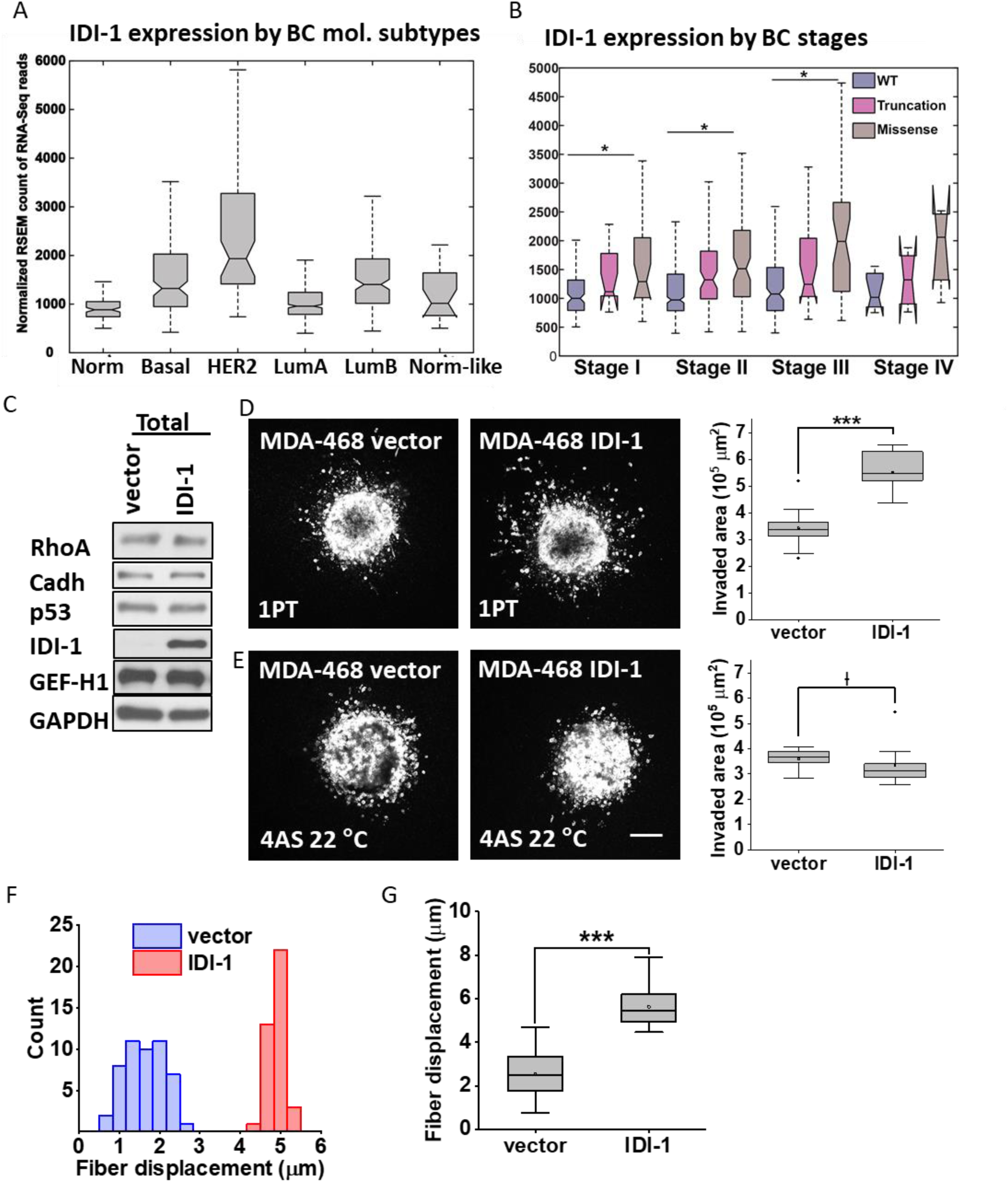
Upregulation of MVA-pathway enzyme IDI-1 expression is sufficient to drive cancer cell invasion in 3D. A) Transcript expression of IDI-1 in TCGA breast cancer dataset (from processed RNA-seq data) stratified based on molecular subtype of the cancer. B) Transcript expression of IDI-1 in TCGA breast cancer dataset (from processed RNA-seq data) stratified based the stage of the cancer and TP53 mutation status. C) Western blot analysis of IDI-1, p53, RhoA, Cadherin and GEF-H1 expression in the MDA-MB-468-IDI-1 stable cell line and the respective vector control cell line. D) Confocal fluorescence maximum projections and invasion quantification of a representative MDA-MB-468-IDI-1 vs -vector control MTS invasion assay in collagen matrix at 24 h. Scale bar = 200 μm. n = 12 and 11 for vector control and IDI-1 cell line, respectively. Statistical significance in this and following invasion assay was determined using Wilcoxon Rank Sum test. E) Confocal fluorescence maximum projections and invasion quantification of a representative MDA-MB-468-IDI-1 vs -vector control MTS invasion assay in a bundled collagen matrix at 24 h (MTSs from the same biological replicate as shown in D). Scale bar = 200 μm. n = 10 (MTSs from the same biological replicate as used in D). F, G) Quantitative analysis of cell-mediated collagen fiber re-organization at the edge of an individual MTS prepared from MDA-MB-468-vector control and MDA-MB-468-IDI-1 cells as assessed through displacement of collagen-embedded fluorescent beads. Bead displacements after 2 hours were quantified for one experiment (F) and for 3 biological replicates (G). n (total number of beads per condition) = 117 and 128 for vector and IDI-1, respectively. Statistical significance was determined via Kolmogorov-Smirnov test.

This observation prompted us to analyze if high expression of the IDI-1 enzyme is sufficient to drive cancer cell invasion in our experimental system. To this end we generated MDA-MB-468 cell lines stably expressing IDI-1 or the respective vector control and confirmed by Western blot analysis that IDI-1 overexpression does not impact p53, RhoA or GEF-H1 levels (Fig. 7c). Subjecting these cells to the MTS invasion assay in low density collagen I matrices revealed a nearly two-fold increase in the mean invaded area in IDI-1 expressing cells in comparison to the vector control cell line indicating that high expression of the IDI-1 protein is sufficient to significantly enhance 3D invasion of cancer cells (Fig. 7d; Fig. S9e, g). Transient knockdown of IDI-1 from these cells confirmed that IDI-1 expression was necessary and mediating the enhanced invasiveness in these cells (Fig. S9i,j). Interestingly, this invasive advantage was abolished during invasion in bundled collagen (Fig. 7e; Fig. S9f, h) suggesting that high expression of IDI-1 supports 3D invasion under conditions that require efficient Rho/ROCK signaling.

Taken together we have provided evidence that the dependence on mutant p53 for Rho/ROCK signaling requires upregulation of MVA pathway gene expression and that one of the MVA pathway targets which is involved in protein prenylation, IDI-1, when overexpressed can phenocopy the ability of mutant p53 to stimulate MTS invasion. Importantly, the supporting effect of mutant p53 on cancer cell invasion as well as the effect of upregulated MVA pathway depends on the biomechanical conditions of the extracellular matrix during the process of invasion, indicating the crucial role of non-cell-autonomous factors in dictating the contribution of mutant p53 in the process of cell locomotion.

## Discussion

The overwhelmingly high prevalence of TP53 alterations in cancers, either by inactivation, loss or various types of mutations, is well established ^54^. Apart from loss of normal functions, many of the mutations confer the p53 protein with new GOF abilities that provide cancer cells with activities affecting key biological processes associated with cancer cell proliferation and invasion, immunogenicity and metabolic re-wiring ^8,55,56^. In that regard, our work has elucidated the GOF activity of three p53 missense mutants in cancer cell invasion in complex three-dimensional environments and linked it to the lipogenic activity of the mevalonate pathway (MVA) that we previously showed to be enhanced in presence of mutant p53 ^25^. The concept of mutant p53 GOF is apparently not universal and seems to be highly context dependent. For example, Tang et al. showed no difference in colon cancer related metastasis when comparing p53-null and a hotspot mutant p53 tumors ^57^, and more recently it was reported that human breast cancer cell lines harboring mutant p53 displayed no alterations in growth in 2D culture, organoids or metastasis *in vivo* when mutant p53 was knocked out ^58^. On the other hand, there is a wealth of evidence that mutant p53 facilitates metastasis in mouse models as reviewed in ^12^. We propose that the best way to reconcile different conclusions about the modes by which mutant p53 proteins do or do not facilitate processes related to metastasis is to assume that the context of the experimental approach including tumor cell types, mutant alleles and experimental protocols may be of critical importance.

Among the studies that do argue for a GOF activity of mutant p53 there were several reports demonstrating that mutant p53 supports cell locomotion by promoting EMT ^59^, by regulating membrane translocation or recycling of cytokine and integrin receptors ^22,60^, or by favoring an active state of Rac1 ^60^. Most of these studies utilized conventional 2D culture methods and assessed cancer cell invasiveness by analysis of cell locomotion on flat uniform surfaces (wound scratch assays) or transmigration of ECM-coated membranes (Boyden chamber assay) towards certain chemo-attractants. Even though the observed phenotype may be the same in 2D and 3D settings, i.e. mutant p53 supports cell migration, the underlying mechanism is likely different as 2D and 3D cell locomotion differ substantially in their cellular and molecular mechanisms ^61^. To identify the cellular processes that are physiologically relevant to the metastatic disease it is imperative to use experimental models that better approximate the *in vivo* setting in terms of multicellular tumor architecture and the surrounding ECM. In this study we utilized multicellular tumor spheroids (MTS) each comprised of several thousand breast or colon carcinoma cells fully embedded into invasion-supporting 3D collagen I environments as a physiologically relevant model for solid cancer dissemination.

The question we addressed was which of the many pathways that are known to be affected by mutant p53 mediates its pro-invasive activity specifically during the process of 3D dissemination. Our study demonstrates that the mutant p53-dependent invasive advantage during 3D invasion of collagen-rich environments is mediated by Rho/ROCK-dependent cellular mechanisms, as p53 depletion dramatically reduced sensitivity to invasion reduction through pharmacological Rho/ROCK inhibition in all tested cell lines. Moreover, we found that cell contractile forces as assessed in collagen gel contraction assays are significantly reduced upon p53 depletion in both breast (MDA-MB-231 and MDA-MB-468) and colon cancer (HCT116) cells. Importantly, cell-mediated alignment of collagen fibers in the vicinity of an individual spheroid before the onset of invasion was also heavily compromised in mutant p53-depleted MDA-MB-468 spheroids as demonstrated by the approximate two-fold reduction of collagen fiber displacement during pre-invasive activity in spheroid invasion assays. The structural remodeling of the ECM is a consequence of Rho-dependent cell-ECM interactions and has been reported to promote breast cancer invasion in vitro as well as *in vivo* in mouse models ^35,62–65^. Note that in human cancers collagen alignment at tumor borders had been described even before the onset of local dissemination and is a prognostic factor for poor breast cancer survival ^66^. Our data indicates that mutant p53 supports 3D cancer cell dissemination by enhancing collagen fiber reorganization and radial alignment at the onset of invasion thereby providing contact guidance and contributing to directional locomotion of cancer cells in a complex 3D environment. This hypothesis is further strengthened by the result that the pro-invasive effect of mutant p53 is fully abolished in bundled collagen matrices where cell-mediated fiber alignment is minimized due to the stiffness of and bundled collagen architecture in the hydrogel ^39^.

Another invasion-regulating cellular characteristic that is strongly dependent on Rho/ROCK and the balance between Rho, Rac and Cdc42 signaling is cell morphology and the establishment of invasion-mediating cell protrusions ^67^. Rho-dependent cell contractility mediates round cell morphology but also formation of membrane blebs – highly dynamic spherical cell protrusions that have been found to contribute to plasticity and efficiency of cancer cell locomotion in 3D environments ^68–70^. In a previous study we demonstrated that membrane blebs, which were previously postulated to only mediate integrin-independent motility, facilitate a novel mode of cancer cell motility in 3D collagen and constitute cell-ECM interaction hubs, where formation of ECM adhesomes and collagen reorganization occurs ^33^. Here, we show that depletion of p53 in invading MDA-MB-231 spheroids diminishes the proportion of the round cell population and the percentage of bleb-bearing cells. The reduced presentation of clustered membrane blebs detected in our experimental setting likely contributes to the diminished invasive capacity of cells relying on a combination of migratory strategies, such as the cell types used for this study. Thus we postulate that the 3D invasion-supporting activity of mutant p53 is mediated via enhanced Rho/ROCK signaling-mediated remodeling of the collagen microenvironment prior to and establishment of membrane blebs during the process of 3D dissemination.

Rho signaling underlies a dynamic spatiotemporal regulation by various upstream effectors (e.g. integrin receptors), the active/inactive conformation-regulating guanine nucleotide exchange factors (GEFs) and GTPase activating proteins (GAPs), as well as by subcellular localization ^71,72^. Muller et al. reported that mutant p53 regulates signaling by α5β1 integrin ^22^. We addressed if β1 integrin protein levels or subcellular distribution are regulated by mutant p53 in our experimental system. FACS analysis of cell surface expression of β1 integrin demonstrated that integrin β1 cell surface presentation was not diminished in p53-depleted cells in any of the cell lines used in this study. The phosphorylation of FAK, one of the most upstream and most prominent components of integrin signaling, as well as the phosphorylation of AKT, were also unaffected by p53 knockdown. These results indicate that, under our experimental setting, the observed reduction in RhoA-dependent cellular processes is not caused by a perturbation in β1 integrin signaling. Another reported p53-dependent regulatory mechanism of RhoA activity is expression of GEF-H1, a guanine exchange factor-H1 for RhoA that is transcriptionally regulated by the V157F, R175H, and R248Q p53 mutants in the osteosarcoma U2OS cell line ^73^. We did not observe an alteration of GEF-H1 expression upon p53 knockdown in MDA-MB-468 breast cancer cells as shown by Western blot protein level analysis (Fig. 5f). Thus, it is not RhoA activation downstream of integrin receptors or GEF-H1 abundance that cause the p53-dependent regulation of Rho/ROCK-dependent cellular processes. Rather, in our experimental conditions, it is the RhoA localization to plasma membrane, a process required for efficient effector binding and downstream activation, that is compromised under depletion of mutant p53. Our data show significantly reduced membrane localization of RhoA as demonstrated by cell fractionation and confocal imaging of immunolabeled endogenous RhoA in collagen-invading spheroids (Fig. 5). This is in agreement with previous reports that mutant p53 affects subcellular distribution of proteins that must be prenylated for correct membrane localization through transcriptional regulation of the mevalonate pathway and of enzymes involved in the process of the lipid moiety transfer ^74^. Here, we report that the invasive capacity of p53-depleted cancer cells used in this study can be fully rescued to the level of p53-bearing cells by exogenous geranylgeranylpyrophosphate (GGPP) supplementation during invasion. Interestingly, the invasive capacity of wildtype p53-bearing HCT116 cells can be increased to the level of mutant p53-expressing HCT116 cells by GGPP addition. Moreover, overexpression of IDI-1, an enzyme contributing to production of the GGPP moiety, is sufficient to significantly increase MDA-MB-468 invasion, thus supporting our hypothesis that it is protein prenylation that mediates the GOF activity of mutant p53 during 3D cancer cell invasion. Here we have provided evidence that mutant p53 plays key roles in the initial steps of egress of cancer cells from a primary tumor and their dissemination in collagen-rich environments. Future studies will hopefully elucidate in which particular scenarios this role is crucial and presents a vulnerability that can be targeted for therapeutic interventions.

## Materials and Methods

### Cell lines

MDA-MB-231 harboring mutant p53 (R280K) and MDA-MB-468 harboring mutant p53 (R273H) breast cancer cells were obtained from the American Type Culture Collection (ATCC) (Manassas, VA). Stable MDA-MB-231 p53 Knock-out (KO) and control parental cell lines were generated using CRISPR-Cas9 editing as described previously ^75^.

The HCT116 parental colorectal adenocarcinoma cell line with wild-type p53 was a gift from Bert Vogelstein. These cells were used to generate p53 knock-out (KO) cells as described ^75^. To generate p53 KO clones, cells were transfected with a pool of three plasmids co-expressing Cas9 and single-guide RNA (sgRNA) targeting p53 exons 4, 5, and 7. Two days later, cells were treated with 10 μM Nutlin-3 for 14 days to inhibit proliferation of cells with wild type p53, thereby enriching for p53 knockout cells as previously described ^76^. Loss of p53 expression in the KO cell line was verified by Western blotting.

The HCT116 cell line expressing p53 R282W hotspot mutant was engineered using base editing technology. An optimized version of the cytosine deaminase base editor11 (a kind gift from Prof. Lukas Dow) was transfected into cells along with a plasmid expressing sgRNA targeting a region of p53 with the desired cytosine-to-thymine mutation in the 4-to 8-base pair window of the protospacer. Single-cell clones were isolated via limiting dilution, clones containing the desired mutation were distinguished via genomic DNA and cDNA sequencing and pooled together.

The MDA-MB-468 cell line expressing IDI-1 was generated as following. The human IDI-1 (RC214341) gene was purchased from OriGene Technologies and cloned into pMXs-IRES-puro retroviral expression vector (Cell Biolabs). The retroviral expression vector and empty vector were transfected into HEK293T cells with packaging plasmids (pCMV-gag-pol and pMD2.G) to produce viruses. MDA-MB-468 cells were infected with retroviruses in the presence of 8 μg/ml polybrene (EMD Millipore). Two days after infection, cells were selected with puromycin at 1 μg/ml for at least 7 days to establish the stable cell lines. The following cloning primers were used:

IDI1-5p-BamHI TAAGGATCCatgtggcgtggactggcgctg

IDI1-3p-XhoI TTACTCGAGtcacattctgtatattttc

### Reagents

Accutase was purchased from MP Biomedicals (Solon, OH). Corning Cell Recovery Solution was purchased from Thermo Fisher Scientific (Waltham, MA). Pepsin-treated (PT) bovine type I collagen was obtained from Advanced BioMatrix (cat.#5010, San Diego, CA) as a 5.9-6.1 mg/ml solution. Acid-solubilized (AS) rat tail type I collagen was obtained as a 10 mg/ml solution from Corning (cat.#354249, Corning, NY). Growth factor-reduced, Phenol Red-free basement membrane extract (BME)/Matrigel was obtained as an 8.9-10 mg/ml solution from BD Biosciences (San Jose, CA). Dimethyl sulfoxide (DMSO) and acetic acid (99.7%) were purchased from Sigma Aldrich (St Louis, MO). DMEM solution (10×), NaOH (1 N) and sodium bicarbonate solution (7.5%) were purchased from Sigma Aldrich and sterile filtered before use. Gibco 4-(2-hydroxyethyl)-1-piperazineethanesulfonic acid (HEPES) buffer (1 M) was obtained from Invitrogen (Carlsbad, CA) and sterile filtered before use. MBQ-167 (inhibition of Rac1/2/3 and Cdc42, ^77^) and Y27362 (inhibition of ROCK, ^78^) inhibitors were obtained from Selleck Chemicals (Houston, TX). Buffered formalin phosphate (10%) was obtained from Fisher Scientific (Pittsburgh, PA). Bovine Serum Albumine (BSA) and Triton-X were obtained from Sigma Aldrich (St Louis, MO).

#### Constructs

Silencer Select Negative Control siRNA #1 (termed “siC” throughout the manuscript, cat.#4390843) and ON-TARGETplus Non-targeting Pool (termed “spC”, cat.# D-001810-10-05) were used as transfection control for Silencer Select siRNAs and SmartPool siRNAs, respectively. For p53 knockdown Silencer Select s605 (termed si#1 in the manuscript, cat.#4390824) and ON-TARGETplus Human TP53 SmartPool (termed sp p53 in the manuscript, cat.# L-003329-00-0005) siRNAs were used. Silencer select siRNAs, Lipofectamine RNAiMax and Opti-MEM were obtained from Thermo Fisher Scientific (Waltham, MA). ON-TARGETplus non-targeting (spC) and TP53-targeting SMARTpool (sp-p53) siRNAs were obtained from Horizon Discovery (Lafayette, CO). DharmaFect1 transfection reagent was obtained from Fisher Scientific (Pittsburth, PA).

#### Antibodies

Antibodies against N-cadherin (cat.#13116S), integrin β1 (cat.#9699), cofilin-1 (cat.#5175T) and phospho cofilin-1 (Ser3) (cat.#3313T) were purchased from Cell Signaling Technology. Rabbit anti-KI67 (ab16667) was purchased from Abcam. Anti-β-actin (cat.#A2066 and A2228), anti-mouse peroxidase (cat.#A4416) and anti-rabbit peroxidase (cat.#A6154) were purchased from Sigma-Aldrich. Anti-E-cadherin (cat.#sc-8426) and unconjugated anti-RhoA (cat.#sc-418) for WB detection were purchased from Santa Cruz Biotechnology. Anti-p53 DO-I and 1801 monoclonal antibodies were purified from hybridomas produced in-house. Rabbit anti-integrin β1 antibody for detection of endogenous β1 integrin on SDS-PAGE was obtained from Cell Signaling (clone D2E5, Danvers, MA), rabbit anti-GAPDH was obtained from Abcam (Cambridge, MA) and horseradish peroxidase (HRP)-conjugated anti-rabbit and anti-mouse IgG antibodies were obtained from Sigma Aldrich (St Louis, MO). Primary and secondary antibodies for immunoblotting were used at 1:1000 and 1:2000 dilution, respectively, unless otherwise noted.

AlexaFluor488-conjugated anti-integrin β1 antibody (clone P5D2, staining total β1) was obtained from Abcam (Cambridge, MA) and used for immunofluorescence, AlexaFluor488-conjugated anti-integrin β1 integrin (clone TS2/16, staining total β1) was obtained from Biolegend (cat.#303007) and used for FACS analysis. AlexaFluor594-conjugated RhoA antibody (clone 26C4) was obtained from Santa Cruz (Dallas, TX).

### Cell culture and analysis

Cell lines were cultured in growth medium consisting of 1× high glucose DMEM containing 10 % (v/v) heat-inactivated fetal bovine serum (FBS) (cat.#900108H, Gemini), 100 units of penicillin, 100 mg/ml streptomycin (cat.#15140122, Thermo Scientific) and 1 % (v/v) 100× non-essential amino acids solution. Cells were maintained at 37 °C with 5 % CO_2_ and were sub-cultured using trypsin-EDTA (cat.#15090-046, Gibco) at 80-90 % confluency. For cell detachment prior to an experiment Accutase was used. All cells used in this study have been tested for mycoplasma contamination and confirmed to be mycoplasma-free. All experiments have been performed with cells between passages 6 to 12 for parental cell lines acquired from ATCC and passages 2 to 8 for stable cell lines with passage 0 being the start of cell culture from a cryopreserved aliquot of cells that underwent the antibiotic selection or the CRISPR procedure. This is especially important for invasion assays which we found sensitive to cell culture conditions and passage number.

#### Transfection of siRNA

For transient siRNA transfection adherent cells were seeded at 6*10^4^ or 8*10^4^ per cm^2^ 24 h prior to transfection and transfected using Lipofectamine RNAiMax transfection reagent (cat.#13778150, Thermo Scientific) or DharmaFect1 transfection reagent (Horizon, cat.# T-2001-03) according to the manufacturer’s instructions.

#### Protein extraction and Western Blotting

Cells were harvested with Accutase or scraped on ice in ice-cold PBS. Cell pellets were lysed with RIPA buffer (50 mM TrisHCl at pH 7.4, 120 mM NaCl, 0.25 % w/v sodium deoxycholate, 1 % v/v Nonidet P-40 with 0.5 mM PMSF, and inhibitor cocktail containing 100 µM benzamidine, 300 µg/µL leupeptin, 100 mg/mL bacitracin, 1 mg/mL a2-macroglobulin. Cell lysates were cleared by spinning at 4000 rpm for 15 min at 4 °C. Protein concentrations were determined using the Bradford assay (Bio-Rad protein assay, Life Science Research). Equivalent amounts of each clarified cell lysate were supplemented with protein sample buffer and incubated for 10 min at 95 °C for denaturation, after which samples were loaded onto a 10-14 % polyacrylamide gel and separated using constant voltage. Proteins were transferred to nitrocellulose (Bio-Rad) or PVDF membranes (IPVH00010, Immobilon, Millipore), blocked for 1 h at room temperature (RT) with PBS containing 0.1 % Tween 20 (Sigma-Aldrich) and 3 % w/v BSA, and probed with the indicated antibodies at 4 °C overnight. Membranes were washed with TBS supplemented with 0.1 % Tween20 prior to the addition of secondary antibodies. Secondary horseradish peroxidase (HRP)-conjugated goat-anti-mouse or goat-anti-rabbit antibody (Sigma) was added and incubated for 1 h at RT. After three washes, immunoblots were visualized by chemiluminescence detection using chemiluminescent horseradish peroxidase reagents (ECL Western Blotting Substrate from Thermo Fisher, Pierce, cat# 32106 or Immobilon Western Chemiluminscent HRP substrate from Millipore, cat# WBKLS0050) according to the manufacturer’s instructions. Densitometry-based quantification of protein expression was performed using ImageJ/Fiji software.

#### Pharmacological cell treatments

Inhibition of mevalonate pathway was achieved by treatment with 10 μM Simvastatin, Fatostatin or GGTI-2133. Fully formed spheroids were pre-treated with the inhibitor or respective solvent control for 4 h at 37 °C in liquid culture, before they were embedded in collagen gels with the inhibitor present at the same concentration both in the gel and the overlaying media for the entire duration of the invasion assay, i.e. 24 h.

Inhibition of Rho signaling was achieved using 15 µM Rhosin, an inhibitor of RhoA, or 10 µM Y27632, an inhibitor of ROCK. Each inhibitor was present at the respective concentration both in the collagen gel and the overlaying media for the entire duration of the invasion assay. Inhibition of Rac/Cdc42 was achieved using 500 nM MBQ-167. All inhibitors were present at the given concentration in the collagen gel and the overlaying media for the entire duration of the invasion assay.

#### FACS analysis of integrin receptor expression

For quantification of cell surface β1 integrin expression 6*10^5^ cells were seeded in 6-well plates and in case of MDA-MB-468 cells transient transfection with p53-targeting or control siRNAs was performed as described above. After 48 h cells were detached and stained using AlexaFluor-conjugated antibody against β1 integrin (clone TS2/16, Biolegend, cat#303007). Briefly, cells were detached using 5 mM EDTA and washed with FACS buffer (HBSS supplemented with 1 % BSA and 0.1 % v/v sodium azide). Samples were then blocked using 10 % Normal Goat Serum (Invitrogen, cat# 50062Z) followed by incubation with the conjugated antibody. Next, samples were washed twice with FACS buffer and resuspended in the same buffer for analysis by flow cytometry in a FACS Celesta, using FACS diva software (BD). All staining and washing procedures were performed at 4 °C.

#### Cell fractionation

Cell fractionation was performed using the Pierce Mem-PER Plus Membrane Protein Extraction Kit from Pierce (Fischer Scientific). In brief, at 72 h post transient transfection, cells were detached using Accutase and 5*10^6^ cells were seeded on collagen I-coated 10 cm culture dishes. Cells were incubated for 90 min at 37 °C and then used for fractionation following the manufacturer’s instructions.

### Preparation and characterization of collagen I-based hydrogels

#### Preparation of cell-free gels

Collagen gel solutions from 1.0 to 4.0 mg/ml were prepared by diluting the high-concentration collagen stock solutions. Appropriate amounts of collagen stock solution were prepared with 10 % (v/v) 10× DMEM, 2.5 % (v/v) HEPES buffer, 2.5 % (v/v) sodium bicarbonate and distilled, deionized, sterile-filtered water. To prevent self-assembly of collagen monomers, all solutions were held and mixed at 4 °C. NaOH was added to adjust the pH to 7.4, and the gel was transferred immediately to the chosen gelation temperature (22 °C or 37 °C). Collagen solutions were allowed to gel for 45 min, which was sufficient to complete gelation at either temperature, and then transferred to an incubator at 37 °C.

To prepare collagen/BME composite gels, first 10× DMEM, HEPES buffer, and sodium bicarbonate were mixed. Then, the required amount of BME stock solution was added. This solution replaced a proportion of the H2O that would be added in the equivalent pure collagen gel. Subsequently the collagen stock solution was added and the solution was brought to pH 7.4 by adding NaOH. After careful mixing, the solution was transferred to the chamber and gelled at 37 °C.

#### Fluorescent Labeling of Collagen

Collagen monomers were labeled with AlexaFluor 647 NHS ester (Thermo Fisher Scientific cat.#A20006) using the method previously described ^79,80^. The labelling procedure was followed for both PT collagen (Advanced Biomatrix) and for AS collagen (Corning). In brief, the collagen stock solution (Advanced Biomatrix 5010) was diluted with sodium bicarbonate to bring the pH to approximately 8.5. Collagen-fluorophore solution was left overnight to react at 4 °C. The resulting collagen solution was re-solubilized and unreacted fluorophore removed by dialyzing against 20 mM acetic acid for 3 days at 4 °C.

#### Collagen Imaging

To prepare collagen gels for imaging, a stock of AlexaFluor 647-labelled collagen was diluted with unlabeled collagen to the desired concentration. For composite collagen/BME gels, HiLyte 488-labelled Laminin (Cytoskeleton Inc.) was mixed with regular BME to a proportion of 2 % by wt. labelled laminin and used in addition to the labelled collagen stock. The gels were prepared in a glass bottom dish following addition of 10x and neutralization with sodium hydroxide. Imaging was performed on a Zeiss LSM800 inverted confocal laser scanning microscope with a 640 nm excitation laser using a 63x oil (NA 1.4) objective in Airyscan mode for the bundled collagen (4 mg/ml AS collagen gelled at 22 °C), and on a Zeiss Elyra 7 with 488 nm and 642 nm excitation lasers using a 63x water (NA 1.2) objective in Lattice SIM mode for low density (1 mg/ml PT collagen gelled at 37 °C), high density (3 mg/ml PT collagen gelled at 37 °C) and mixed collagen ( 1mg/ml PT collagen with 3 mg/ml BME gelled at 37 °C) gels.

#### Pore Size Determination

Pore sizes were determined using the method described in ^81^. Single slices of collagen gel images were pre-processed in ImageJ before to yield a segmented image that was then used for determination of the pore size. Raw images were subjected to a bandpass filter and rolling ball background subtraction to remove features below a certain size threshold and reduce out of plane background signal, respectively. Images were then subjected to an auto local threshold to obtain a binarized image of the network. To determine pore size, the distances between “on” pixels were counted row by row and column by column in the segmented image. The distribution of these gaps between fibers was fit to the exponential probability density function:

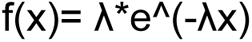

The pore size reported for the network is determined by the characteristic pore size 1/ƛ.

#### Rheology

Rheological testing of gels was performed on an Anton-Paar MCR302 rheometer with a 50-mm polycarbonate cone-and-plate geometry (1° angle and 0.097 mm truncation) as the top plate and a glass bottom plate. The temperature (22, 25 or 37 °C) of the bottom plate was preset and maintained throughout measurement using a Peltier control unit. A solvent trap and sample hood was used to minimize evaporation. 700 µl of gelling solution were placed upon the bottom plate and the measuring geometry was brought to the fixed 0.097 mm gap. Any excess solution remaining after the top plate was at the measuring distance was carefully wiped away to avoid overfilling. Storage modulus and loss modulus were then measured in strain controlled oscillatory shear with 1 % strain and 1 Hz oscillations.

#### Preparation of cell-embedded gels

To prepare collagen gels loaded with a single spheroid, 1 mg/ml collagen solution was prepared as described above at 4 °C to prevent gelation. Neutralized collagen solution (150 µl) was added to a well of a flat bottom 96-well plate or a chamber consisting of a 5 mm glass cylinder glued to a coverslip-bottom cell culture dish. Spheroids were individually added to the liquid collagen in 5 µl liquid. The plate was then transferred immediately to a 37 °C incubator and after 1 h the collagen gels were overlaid with 40 µl growth medium. To prepare collagen/BME composite matrices loaded with a single spheroid, the wells were first pre-coated with 40 µl collagen/BME solution to prevent sedimentation of spheroids to the bottom of the chamber. The coating was gelled for 1 h at 37 °C. Following this, the bulk volume of the matrix solution was added to the chamber and a single spheroid was implanted as described above.

To prepare collagen gels loaded with dispersed cells, the collagen solution was prepared omitting part of the water and neutralized at 4 °C. The water was substituted with growth medium containing the desired number of cells. Cell-loaded collagen was gelled at 37 °C as described above for MTS-loaded collagen gels.

### Generation and characterization of multicellular tumor spheroids (MTS)

Spheroids were formed from 3000 (for invasion assays) to 10000 cells/spheroid (for cryosectioning of invading spheroids) in appropriate growth medium supplemented with 0.25 mg/ml BME using a centrifugation method described previously ^29^ employing ultralow attachment 96-well U-bottom Nunclon Sphera plates from Thermo Fisher Scientific (Waltham, MA). HCT116 spheroids were formed in BME-free growth media using the same centrifugation method. Spheroids were allowed to form for 48 h at 37 °C under 5 % CO_2_.

#### 3D Cell Proliferation assay

To compare cell proliferation and viability in 3D cell cultures, the Cell Titer Glo 3D assay (Promega) was used according to manufacturer’s instructions. In brief, for each biological replicate individual spheroids were transferred in 100 µl growth media to a new 96-well plate, mixed with 100 µl pre-warmed Cell Titer Glo 3D reagent and shaken for 20 minutes at room temperature. Subsequent reading was performed using standard settings in a Synergy H1 Hybrid plate reader (Biotek). 6 spheroids per condition were used for each independent biological replicate.

#### Spheroid invasion assay

Fully formed spheroids prepared from cells labelled with a cytosolic dye were transferred to Cell Recovery Solution (200 μl per spheroid) and incubated 45 -60 min at 4 °C to remove any remains of BME used during spheroid formation. Subsequently, spheroids were embedded into 150 μl collagen I or composite collagen I/BME gels and transferred to 37 °C for gelation of the biomatrix unless stated otherwise. After completion of gelation (1 h) gels were overlayed with 40 μl of growth media and imaged at 10x magnification on a transmitted light microscope in the DIC setting to capture the initial size of each spheroid before the onset of invasion (t=0). For analysis of invasion distance, spheroids were imaged at 10x magnification in confocal fluorescence mode at 24 h after implantation. For each spheroid, we used these magnified images to determine the invaded area, defined as the difference between the initial area of the spheroid at t0 and an ellipse that circumscribes 90% of the invasive cells at t = 24 h. In Figure Legends describing graphical data with spheroids, n signifies the number of spheroids per experimental condition, unless stated otherwise.

#### Immunocytochemistry of spheroids

For immunocytochemical staining fully formed and CRS-treated MTSs were embedded in 1 mg/ml collagen matrices of 100-150 μl. Samples were fixed in neutrally buffered pre-warmed to 37 °C 4 % formalin solution for 15 min at 24 °C. After extensive washing and permeabilization with 0.2 % Triton-X, samples were washed again with PBS to remove the detergent and samples were blocked with 1 % BSA/PBS for 1 h at 24 °C. Subsequently the blocking solution was removed and fluorescently labeled phalloidin (1:1000), DAPI (1:1000 for cryosections,1:500 for 3D) and, if needed, the respective directly labeled antibody (anti-RhoA 1:100 for cryosections, anti-β1 integrin 1:500 for cryosections, 1:300 for 3D) were added and the samples were incubated for 16-24 h at 4 °C. After extensive washing with PBS, samples were overlaid with PBS and immediately subjected to imaging.

#### Collagen contraction assay

Collagen solution containing 1*10^6^ cells/ml were prepared, pre-nucleated for 10 min at room temperature and 600 μl gels were cast onto the 23 mm coverslip bottom of pre-warmed FluroDishes (35 mm). Gels were polymerized at 37 °C, overlaid with 1.5 ml growth medium, and manually released from the glass bottom. Contraction was allowed for 24 h at normal incubation conditions. Gel contraction was documented using digital photography. Images were taken at t = 0 h (before gel release) and at t = 24 h, and gel area was measured. Contraction was expressed as a percentage decrease of gel area over 24 h. All conditions were tested with three or four technical replicates.

#### Spheroid Contraction and SIM Imaging

For contraction assays spheroids were prepared as described and after 48 hours were deshelled for 40-45 min in Cell Recovery Solution (Corning, 354253). Spheroids were then implanted into 1 mg/mL PT collagen gels which were prepared as described in section “Preparation of cell free gels”. In addition to the typical gel solution components, fluorescently labeled AlexaFluor647-PT collagen monomer (10 % w/w relative to unlabeled collagen) and 1.0 µm polystyrene fluorescent beads (1.3 % v/v from stock solution, Thermo Fischer 13083) were added for visualization of collagen fiber and overall gel displacements, respectively. Immediately after casting the gel, the spheroid laden gel was moved to an incubated microscope stage (kept at 37 °C and 5 % CO_2_ for the duration of the experiment) and allowed to gel for 20 min, just until stable collagen fibers had formed. The initial positions of fibers and beads were recorded as this time and then again at 120 minutes using structured illumination microscopy (SIM).

### Image acquisition and analysis

For lung tissue colonization assay, brightfield images of H&E stained samples were taken with Leica AT2 whole-slide digital imaging (Leica).

For MTS invasion analysis, imaging was performed on an inverted confocal laser-scanning microscope (Zeiss LSM 800) in confocal fluorescence mode with 10x air (NA 0.4). An Airyscan system was used for detection on the Zeiss LSM 800. Confocal fluorescence z-stacks (step size 10 µm) of fluorescently labeled spheroids were acquired at 1 h and 24 h after implantation into the collagen matrix. We used maximum intensity projections of these magnified fluorescence images to determine the invaded area, defined as the difference between the area of the spheroid at t = 1 h and an ellipse that circumscribes 90% of the invasive cells at t = 24 h.

For analysis of cell morphology and protrusions, confocal fluorescence z-stacks (step size 1 µm) of spheroids fixed after 48 h of invasion in collagen matrices and stained with fluorescently labeled phalloidin were acquired on a Zeiss LSM 800 using the 20x objective. Cell morphology was determined in a semi-automated fashion using ImageJ. First, maximum fluorescence intensity projections were generated from z-stacks, despeckled and thresholded. Following this processing, the Analyze Particle function was used to assess the circularity, c, of each cell, with 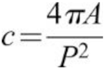, where A is cell area and P the cell perimeter. Circularity thus ranges from 0 to 1, with c = 1 for a circle. Cell protrusion classification was performed by eye using the preprocessed images.

For analysis of single MTS collagen contraction imaging was performed on Zeiss Elyra using a 63x water immersion objective (NA 1.2). The initial positions of fibers and beads were recorded upon appearance of stable collagen fibers (approx. 20 min) using structured illumination microscopy (SIM). The spheroid was allowed to contract the gel for an additional 100 minutes (120 min total time in gel), after which a final SIM image was recorded. SIM images were post-processed in Zeiss software to yield the reconstructed super-resolution image. Spheroid contraction was quantified by measuring the distance between the initial and final positions of fluorescent beads in SIM images using ImageJ.

### *Ex vivo* colonization of lung tissue

The lung decellularization process and *ex vivo* cultures were modified from a published protocol ^28^. All animal studies were approved by Institutional Animal Care and Use Committee at Columbia University, and all experiments were conducted in compliance with the NIH guidelines for animal research. Briefly adult C57BL/6 mice were euthanized, and an incision was made along the sternum to expose the trachea. The lungs were rinsed and inflated with rinse solution (deionized water + 5x Pen/Strep) that was injected through trachea with a 21-guage needle, and then were excised with intact trachea and heart, placed in a sterile 6-well plate, and incubated in rinse solution at 4 °C for 1 hour. Subsequently, the lungs underwent a series of rinses and incubations on a rotator to remove cellular components and DNA: First, lung tissues were injected with five rinses of 3 mL rinse solution through the trachea. Subsequently, Triton solution (3 mL; 0.1 % Triton X-100 with 5x Pen/Strep) was introduced through the tracheal opening, and the lungs were incubated overnight at 4 °C to lyse cells. Following this incubation, the lungs were taken out of the Triton solution and underwent five additional washes with the rinse solution. Next, the lungs were exposed to deoxycholate solution (3 mL; 2 % sodium deoxycholate; SDC with 1x Pen/Strep) injected through the tracheal aperture, and incubated overnight at 4 °C in an orbital to facilitate the removal of cellular components. A NaCl solution (3 mL; 1 M NaCl with 5x Pen/Strep) was then injected through the same tracheal opening, and the lungs were incubated for 1 hour at room temperature to lyse any remaining nuclei. Finally, a DNase solution (3 mL; 30 mg/mL porcine pancreatic DNase in 1.3 mM MgSO4 and 2 mM CaCl2 with 5x Pen/Strep) was introduced through the tracheal aperture, and the lungs were incubated for 1 hour at room temperature to mediate DNA lysis. The lungs were then removed from the DNase solution and rinsed with a phosphate-buffered saline (PBS) solution.

The resulting decellularized lung tissue was cut into small cubes and incubated in a corresponding media for 2 hours. The lung pieces were placed on a 12-well plate and 5.0*10^5^ MDA-MB 231 cells were seeded on the tissues, incubated at 37 °C for 2 hours to facilitate cell attachment. Unattached cells were gently rinsed with PBS. Each well was filled with 2 ml DMEM + 10 % FBS + 1 % Pen/Strep. The *ex vivo* cultures were incubated at 37 °C with 5 % CO_2_ for 10 days and media was changed once every 2 days. Tissues were fixed with 4 % PFA for 4 hours at 4 °C, embedded in paraffin and sectioned. The sections were stained for Hematoxylin & Eosin (H&E) and brightfield images were acquired. The number of colonized cells were quantified using QuPath software.

For immunohistochemistry staining the decellularized lung tissues were fixed in buffered zinc formalin (Thermo Fisher, 22-050-259) for 18 hours at 4°C and placed in 70% ethanol. Antigen retrieval was performed on FFPE sections using 10mM boiling citric acid buffer (pH=6) (immunohistochemistry) in a pressure cooker. Endogenous peroxidases were quenched with 3% H_2_O_2_ (Sigma Aldrich, H1009), blocked with Avidin (Sigma-Aldrich, A9275), Biotin (Sigma-Aldrich, B4501) and Starting Block T20 buffer (Thermo Fisher, 37543) for 15 minutes each at room temperature. The tissue sections were incubated with rabbit anti-KI-67 primary antibody (Abcam, ab16667) for approximately 16 hours at 4°C, and then incubated with a goat anti-rabbit IgG (H+L) biotinylated secondary antibody (Vector Laboratories, BA-1000-1.5) for 30 minutes at 37°C. The antibodies were diluted in PBS+1% Bovine serum albumin+0.3% TritonX buffer. Then slides were incubated with ABC reagent (Vector Laboratories, PK-6100) for 30 minutes at 37°C, treated with DAB substrate kit (Vector Laboratories, SK4100), counterstained with Hematoxylin, dehydrated and mounted for imaging. We thank the Confocal and Special Microscopy, Molecular Pathology and Biostatistics Shared Resources at the Herbert Irving Comprehensive Cancer Center (HICCC).

### Statistical analysis of experiments and TCGA patient datasets

All statistical tests were conducted at α = 0.05. Statistical significance is indicated by *P<0.05, **P<0.01 and ***P<0.001. Lack of statistical significance (P>0.05) is denoted by a dagger symbol (†).

The number of colonizing cells and KI67^+^ cells in lung tissue colonization assays was compared using Wilcoxon Rank Sum test.

For analysis of spheroid invasion, Wilcoxon Rank Sum tests were used to compare spheroid invasion under genetic depletion or pharmacological inhibition to the appropriate control.

Circularity and cell protrusion distributions were compared using Kolmogorov–Smirnov (KS) analysis.

For the collagen contraction assays, the percentage of contracted collagen area of p53-depleted cells was compared to the p53+/+ cells using two-sample unequal variance t-test (Welch’s t-test).

For quantitative analysis of collagen fiber displacement in the vicinity of an individual MTS, statistical significance was determined by Kolmogorov-Smirnov test.

Comparison of Pearson’s R values for RhoA and the respective membrane marker in p53+/+ versus p53-depleted cells was carried out using Wilcoxon Rank Sum tests.

#### The Cancer Genome Atlas (TCGA) data analysis

TCGA data was downloaded from the TCGA data portal (later from the GDC data portal) as well as cBioportal. Customized scripts and Matlab programs were used to normalize, quantify and calculate differential expression statistics between samples. Copy number alterations data and sub-type classification data were downloaded from UCSC Xena Functional Genomics Explorer (https://xenabrowser.net/). Boxplots were plotted with Matlab (from Mathworks, versions R2018A, R2020B and R2022A). The Welch’s t-test (unequal variances) was used to determine statistical significance. To correct for multiple testing, when necessary the false discovery rate correction (Benjamini and Hochberg procedure) was used, unless indicated otherwise.

#### Survival analysis

Clinical data including survival data were downloaded from the TCGA data portal (later from the GDC data portal). Samples were sorted on the basis of expression of specific genes. Samples were classified into two groups, high versus low, based on the expression of the gene related to the median expression. The median overall survival and statistical significance (p-value) were calculated as described below. Briefly, Kaplan– Meier curves provide a method for estimating the survival curve, and the log rank test provides a statistical comparison of two groups.

## Acknowledgements

We are grateful to members of the Prives and Kaufman labs for helpful discussions and advice. Ella Freulich is thanked for expert technical assistance in preparing laboratory reagents and cell culture. This work was supported in part by National Institutes of Health (NIH) grants P30CA013696 (C.P, L.J.K and A.K.R.), HICCC Pilot Award (C.P. and L.J.K.), P01-CA098101 (A.K.R., G.E., R.N.), the Ramon Areces Foundation scholarship (R.N.) and the NIH fellowship F31CA275369-02 (G.E.) and R35 CA220526 to CP.

## Bibliography

1 Gupta, G. P. & Massagué, J. Cancer metastasis:: Building a framework. Cell 127, 679–695, doi:10.1016/j.cell.2006.11.001 (2006).

2 Lambert, A. W., Pattabiraman, D. R. & Weinberg, R. A. Emerging Biological Principles of Metastasis. Cell 168, 670–691, doi:10.1016/j.cell.2016.11.037 (2017).

3 Olivier, M., Hollstein, M. & Hainaut, P. TP53 mutations in human cancers: origins, consequences, and clinical use. Cold Spring Harb Perspect Biol 2, a001008, doi:10.1101/cshperspect.a001008 (2010).

4 Lang, G. A. et al. Gain of function of a p53 hot spot mutation in a mouse model of Li-Fraumeni syndrome. Cell 119, 861–872, doi:10.1016/j.cell.2004.11.006 (2004).

5 Olive, K. P. et al. Mutant p53 gain of function in two mouse models of Li-Fraumeni syndrome. Cell 119, 847–860, doi:10.1016/j.cell.2004.11.004 (2004).

6 Hanel, W. et al. Two hot spot mutant p53 mouse models display differential gain of function in tumorigenesis. Cell Death Differ 20, 898–909, doi:10.1038/cdd.2013.17 (2013).

7 Zhang, Y. et al. Somatic Trp53 mutations differentially drive breast cancer and evolution of metastases. Nat Commun 9, 3953, doi:10.1038/s41467-018-06146-9 (2018).

8 Efe, G. et al. p53 Gain-of-Function Mutation Induces Metastasis via BRD4-Dependent CSF-1 Expression. Cancer Discov 13, 2632–2651, doi:10.1158/2159-8290.CD-23-0601 (2023).

9 Brosh, R. & Rotter, V. When mutants gain new powers: news from the mutant p53 field. Nat Rev Cancer 9, 701–713, doi:10.1038/nrc2693 (2009).

10 Muller, P. A. & Vousden, K. H. p53 mutations in cancer. Nat Cell Biol 15, 2–8, doi:10.1038/ncb2641 (2013).

11 Freed-Pastor, W. A. & Prives, C. Mutant p53: one name, many proteins. Genes Dev 26, 1268–1286, doi:10.1101/gad.190678.112 (2012).

12 Tang, Q., Su, Z., Gu, W. & Rustgi, A. K. Mutant p53 on the Path to Metastasis. Trends Cancer 6, 62–73, doi:10.1016/j.trecan.2019.11.004 (2020).

13 Pawluchin, A. & Galic, M. Moving through a changing world: Single cell migration in 2D vs. 3D. Front Cell Dev Biol 10, 1080995, doi:10.3389/fcell.2022.1080995 (2022).

14 Petrie, R. J. & Yamada, K. M. At the leading edge of three-dimensional cell migration. J Cell Sci 125, 5917–5926, doi:10.1242/jcs.093732 (2012).

15 Ridley, A. J. Rho GTPase signalling in cell migration. Curr Opin Cell Biol 36, 103–112, doi:10.1016/j.ceb.2015.08.005 (2015).

16 Friedl, P. & Alexander, S. Cancer invasion and the microenvironment: plasticity and reciprocity. Cell 147, 992–1009, doi:10.1016/j.cell.2011.11.016 (2011).

17 Orgaz, J. L. & Sanz-Moreno, V. Emerging molecular targets in melanoma invasion and metastasis. Pigment Cell Melanoma Res 26, 39–57, doi:10.1111/pcmr.12041 (2013).

18 van Helvert, S., Storm, C. & Friedl, P. Mechanoreciprocity in cell migration. Nat Cell Biol 20, 8–20, doi:10.1038/s41556-017-0012-0 (2018).

19 Haga, R. B. & Ridley, A. J. Rho GTPases: Regulation and roles in cancer cell biology. Small GTPases 7, 207–221, doi:10.1080/21541248.2016.1232583 (2016).

20 Lawson, C. D. & Ridley, A. J. Rho GTPase signaling complexes in cell migration and invasion. J Cell Biol 217, 447–457, doi:10.1083/jcb.201612069 (2018).

21 Mullen, P. J., Yu, R., Longo, J., Archer, M. C. & Penn, L. Z. The interplay between cell signalling and the mevalonate pathway in cancer. Nat Rev Cancer 16, 718–731, doi:10.1038/nrc.2016.76 (2016).

22 Muller, P. A. et al. Mutant p53 drives invasion by promoting integrin recycling. Cell 139, 1327–1341, doi:10.1016/j.cell.2009.11.026 (2009).

23 Arjonen, A. et al. Mutant p53-associated myosin-X upregulation promotes breast cancer invasion and metastasis. J Clin Invest 124, 1069–1082, doi:10.1172/JCI67280 (2014).

24 Timpson, P. et al. Spatial regulation of RhoA activity during pancreatic cancer cell invasion driven by mutant p53. Cancer Res 71, 747–757, doi:10.1158/0008-5472.CAN-10-2267 (2011).

25 Freed-Pastor, W. A. et al. Mutant p53 disrupts mammary tissue architecture via the mevalonate pathway. Cell 148, 244–258, doi:10.1016/j.cell.2011.12.017 (2012).

26 Sorrentino, G. et al. Metabolic control of YAP and TAZ by the mevalonate pathway. Nat Cell Biol 16, 357–366, doi:10.1038/ncb2936 (2014).

27 Esposito, D. et al. ROCK1 mechano-signaling dependency of human malignancies driven by TEAD/YAP activation. Nat Commun 13, 703, doi:10.1038/s41467-022-28319-3 (2022).

28 Xiong, G., Flynn, T. J., Chen, J., Trinkle, C. & Xu, R. Development of an ex vivo breast cancer lung colonization model utilizing a decellularized lung matrix. Integr Biol (Camb) 7, 1518–1525, doi:10.1039/c5ib00157a (2015).

29 Ivascu, A. & Kubbies, M. Rapid generation of single-tumor spheroids for high-throughput cell function and toxicity analysis. J Biomol Screen 11, 922–932, doi:10.1177/1087057106292763 (2006).

30 Paluch, E. K. & Raz, E. The role and regulation of blebs in cell migration. Curr Opin Cell Biol 25, 582–590, doi:10.1016/j.ceb.2013.05.005 (2013).

31 Cantelli, G. et al. TGF-beta-Induced Transcription Sustains Amoeboid Melanoma Migration and Dissemination. Curr Biol 25, 2899–2914, doi:10.1016/j.cub.2015.09.054 (2015).

32 Orgaz, J. L. et al. Diverse matrix metalloproteinase functions regulate cancer amoeboid migration. Nat Commun 5, 4255, doi:10.1038/ncomms5255 (2014).

33 Guzman, A., Avard, R. C., Devanny, A. J., Kweon, O. S. & Kaufman, L. J. Delineating the role of membrane blebs in a hybrid mode of cancer cell invasion in three-dimensional environments. J Cell Sci 133, doi:10.1242/jcs.236778 (2020).

34 Fackler, O. T. & Grosse, R. Cell motility through plasma membrane blebbing. J Cell Biol 181, 879–884, doi:10.1083/jcb.200802081 (2008).

35 Provenzano, P. P., Inman, D. R., Eliceiri, K. W., Trier, S. M. & Keely, P. J. Contact guidance mediated three-dimensional cell migration is regulated by Rho/ROCK-dependent matrix reorganization. Biophys J 95, 5374–5384, doi:10.1529/biophysj.108.133116 (2008).

36 Shang, X. et al. Rational design of small molecule inhibitors targeting RhoA subfamily Rho GTPases. Chem Biol 19, 699–710, doi:10.1016/j.chembiol.2012.05.009 (2012).

37 Uehata, M. et al. Calcium sensitization of smooth muscle mediated by a Rho-associated protein kinase in hypertension. Nature 389, 990–994, doi:10.1038/40187 (1997).

38 Charras, G. & Sahai, E. Physical influences of the extracellular environment on cell migration. Nat Rev Mol Cell Biol 15, 813–824, doi:10.1038/nrm3897 (2014).

39 Xie, J., Bao, M., Bruekers, S. M. C. & Huck, W. T. S. Collagen Gels with Different Fibrillar Microarchitectures Elicit Different Cellular Responses. ACS Appl Mater Interfaces 9, 19630–19637, doi:10.1021/acsami.7b03883 (2017).

40 Oh, S., Nguyen, Q. D., Chung, K. H. & Lee, H. Bundling of Collagen Fibrils Using Sodium Sulfate for Biomimetic Cell Culturing. ACS Omega 5, 3444–3452, doi:10.1021/acsomega.9b03704 (2020).

41 Rosel, D. et al. Up-regulation of Rho/ROCK signaling in sarcoma cells drives invasion and increased generation of protrusive forces. Mol Cancer Res 6, 1410–1420, doi:10.1158/1541-7786.MCR-07-2174 (2008).

42 Yamamoto, A., Doak, A. E. & Cheung, K. J. Orchestration of Collective Migration and Metastasis by Tumor Cell Clusters. Annu Rev Pathol 18, 231–256, doi:10.1146/annurev-pathmechdis-031521-023557 (2023).

43 Pandya, P., Orgaz, J. L. & Sanz-Moreno, V. Actomyosin contractility and collective migration: may the force be with you. Curr Opin Cell Biol 48, 87–96, doi:10.1016/j.ceb.2017.06.006 (2017).

44 Guzman, A., Ziperstein, M. J. & Kaufman, L. J. The effect of fibrillar matrix architecture on tumor cell invasion of physically challenging environments. Biomaterials 35, 6954–6963, doi:10.1016/j.biomaterials.2014.04.086 (2014).

45 van Unen, J. et al. Plasma membrane restricted RhoGEF activity is sufficient for RhoA-mediated actin polymerization. Sci Rep 5, 14693, doi:10.1038/srep14693 (2015).

46 Budnar, S. et al. Anillin Promotes Cell Contractility by Cyclic Resetting of RhoA Residence Kinetics. Dev Cell 49, 894–906 e812, doi:10.1016/j.devcel.2019.04.031 (2019).

47 Avard, R. C. et al. DISC-3D: dual-hydrogel system enhances optical imaging and enables correlative mass spectrometry imaging of invading multicellular tumor spheroids. Sci Rep 13, 12383, doi:10.1038/s41598-023-38699-1 (2023).

48 Istvan, E. S. & Deisenhofer, J. Structural mechanism for statin inhibition of HMG-CoA reductase. Science 292, 1160–1164, doi:10.1126/science.1059344 (2001).

49 Kamisuki, S. et al. A small molecule that blocks fat synthesis by inhibiting the activation of SREBP. Chem Biol 16, 882–892, doi:10.1016/j.chembiol.2009.07.007 (2009).

50 Xue, L. et al. Targeting SREBP-2-Regulated Mevalonate Metabolism for Cancer Therapy. Front Oncol 10, 1510, doi:10.3389/fonc.2020.01510 (2020).

51 Adnane, J., Bizouarn, F. A., Qian, Y., Hamilton, A. D. & Sebti, S. M. p21(WAF1/CIP1) is upregulated by the geranylgeranyltransferase I inhibitor GGTI-298 through a transforming growth factor beta-and Sp1-responsive element: involvement of the small GTPase rhoA. Mol Cell Biol 18, 6962–6970, doi:10.1128/MCB.18.12.6962 (1998).

52 Moon, S. H. et al. p53 Represses the Mevalonate Pathway to Mediate Tumor Suppression. Cell 176, 564-+, doi:10.1016/j.cell.2018.11.011 (2019).

53 Cancer Genome Atlas, N. Comprehensive molecular portraits of human breast tumours. Nature 490, 61–70, doi:10.1038/nature11412 (2012).

54 Leroy, B., Anderson, M. & Soussi, T. TP53 mutations in human cancer: database reassessment and prospects for the next decade. Hum Mutat 35, 672–688, doi:10.1002/humu.22552 (2014).

55 Alvarado-Ortiz, E. et al. Mutant p53 Gain-of-Function: Role in Cancer Development, Progression, and Therapeutic Approaches. Front Cell Dev Biol 8, 607670, doi:10.3389/fcell.2020.607670 (2020).

56 Bargonetti, J. & Prives, C. Gain-of-function mutant p53: history and speculation. J Mol Cell Biol 11, 605–609, doi:10.1093/jmcb/mjz067 (2019).

57 Tang, J. et al. Trp53 null and R270H mutant alleles have comparable effects in regulating invasion, metastasis, and gene expression in mouse colon tumorigenesis. Lab Invest 99, 1454–1469, doi:10.1038/s41374-019-0269-y (2019).

58 Wang, Z. et al. Loss-of-Function but Not Gain-of-Function Properties of Mutant TP53 Are Critical for the Proliferation, Survival, and Metastasis of a Broad Range of Cancer Cells. Cancer Discov 14, 362–379, doi:10.1158/2159-8290.CD-23-0402 (2024).

59 Dong, P. et al. Mutant p53 gain-of-function induces epithelial-mesenchymal transition through modulation of the miR-130b-ZEB1 axis. Oncogene 32, 3286–3295, doi:10.1038/onc.2012.334 (2013).

60 Yue, X. et al. Gain-of-function mutant p53 activates small GTPase Rac1 through SUMOylation to promote tumor progression. Genes Dev 31, 1641–1654, doi:10.1101/gad.301564.117 (2017).

61 Yamada, K. M. & Sixt, M. Mechanisms of 3D cell migration. Nat Rev Mol Cell Biol 20, 738–752, doi:10.1038/s41580-019-0172-9 (2019).

62 Malik, R., Lelkes, P. I. & Cukierman, E. Biomechanical and biochemical remodeling of stromal extracellular matrix in cancer. Trends Biotechnol 33, 230–236, doi:10.1016/j.tibtech.2015.01.004 (2015).

63 Riching, K. M. et al. 3D collagen alignment limits protrusions to enhance breast cancer cell persistence. Biophys J 107, 2546–2558, doi:10.1016/j.bpj.2014.10.035 (2014).

64 Lyons, T. R. et al. Postpartum mammary gland involution drives progression of ductal carcinoma in situ through collagen and COX-2. Nat Med 17, 1109–1115, doi:10.1038/nm.2416 (2011).

65 Alexander, S., Weigelin, B., Winkler, F. & Friedl, P. Preclinical intravital microscopy of the tumour-stroma interface: invasion, metastasis, and therapy response. Curr Opin Cell Biol 25, 659–671, doi:10.1016/j.ceb.2013.07.001 (2013).

66 Conklin, M. W. et al. Collagen Alignment as a Predictor of Recurrence after Ductal Carcinoma In Situ. Cancer Epidemiol Biomarkers Prev 27, 138–145, doi:10.1158/1055-9965.EPI-17-0720 (2018).

67 Riching, K. M. & Keely, P. J. Rho family GTPases: making it to the third dimension. Int J Biochem Cell Biol 59, 111–115, doi:10.1016/j.biocel.2014.11.007 (2015).

68 Bergert, M., Chandradoss, S. D., Desai, R. A. & Paluch, E. Cell mechanics control rapid transitions between blebs and lamellipodia during migration. Proc Natl Acad Sci U S A 109, 14434–14439, doi:10.1073/pnas.1207968109 (2012).

69 Diz-Munoz, A. et al. Steering cell migration by alternating blebs and actin-rich protrusions. BMC Biol 14, 74, doi:10.1186/s12915-016-0294-x (2016).

70 Friedl, P. & Wolf, K. Plasticity of cell migration: a multiscale tuning model. J Cell Biol 188, 11–19, doi:10.1083/jcb.200909003 (2010).

71 Marjoram, R. J., Lessey, E. C. & Burridge, K. Regulation of RhoA activity by adhesion molecules and mechanotransduction. Curr Mol Med 14, 199–208, doi:10.2174/1566524014666140128104541 (2014).

72 de Seze, J., Gatin, J. & Coppey, M. RhoA regulation in space and time. FEBS Lett 597, 836–849, doi:10.1002/1873-3468.14578 (2023).

73 Mizuarai, S., Yamanaka, K. & Kotani, H. Mutant p53 induces the GEF-H1 oncogene, a guanine nucleotide exchange factor-H1 for RhoA, resulting in accelerated cell proliferation in tumor cells. Cancer Res 66, 6319–6326, doi:10.1158/0008-5472.CAN-05-4629 (2006).

74 Etichetti, C. M. B., Zalazar, E. A., Cocordano, N. & Girardini, J. Beyond the Mevalonate Pathway: Control of Post-Prenylation Processing by Mutant p53. Front Oncol 10, doi:ARTN 59503410.3389/fonc.2020.595034 (2020).

75 Klein, A. M. et al. MDM2, MDMX, and p73 regulate cell-cycle progression in the absence of wild-type p53. Proc Natl Acad Sci U S A 118, doi:10.1073/pnas.2102420118 (2021).

76 Tong, D. R. et al. p53 Frameshift Mutations Couple Loss-of-Function with Unique Neomorphic Activities. Mol Cancer Res 19, 1522–1533, doi:10.1158/1541-7786.MCR-20-0691 (2021).

77 Humphries-Bickley, T. et al. Characterization of a Dual Rac/Cdc42 Inhibitor MBQ-167 in Metastatic Cancer. Mol Cancer Ther 16, 805–818, doi:10.1158/1535-7163.MCT-16-0442 (2017).

78 Ishizaki, T. et al. Pharmacological properties of Y-27632, a specific inhibitor of rho-associated kinases. Mol Pharmacol 57, 976–983 (2000).

79 Kalia, J. & Raines, R. T. Advances in Bioconjugation. Curr Org Chem 14, 138–147, doi:10.2174/138527210790069839 (2010).

80 Stephanopoulos, N. & Francis, M. B. Choosing an effective protein bioconjugation strategy. Nat Chem Biol 7, 876–884, doi:10.1038/nchembio.720 (2011).

81 Kaufman, L. J. et al. Glioma expansion in collagen I matrices: analyzing collagen concentration-dependent growth and motility patterns. Biophys J 89, 635–650, doi:10.1529/biophysj.105.061994 (2005).

