## Supplementary Material for "Mutant p53 regulates cancer cell invasion in complex three-dimensional environments through mevalonate pathway-dependent Rho/ROCK signaling"

Suppl. Fig.1

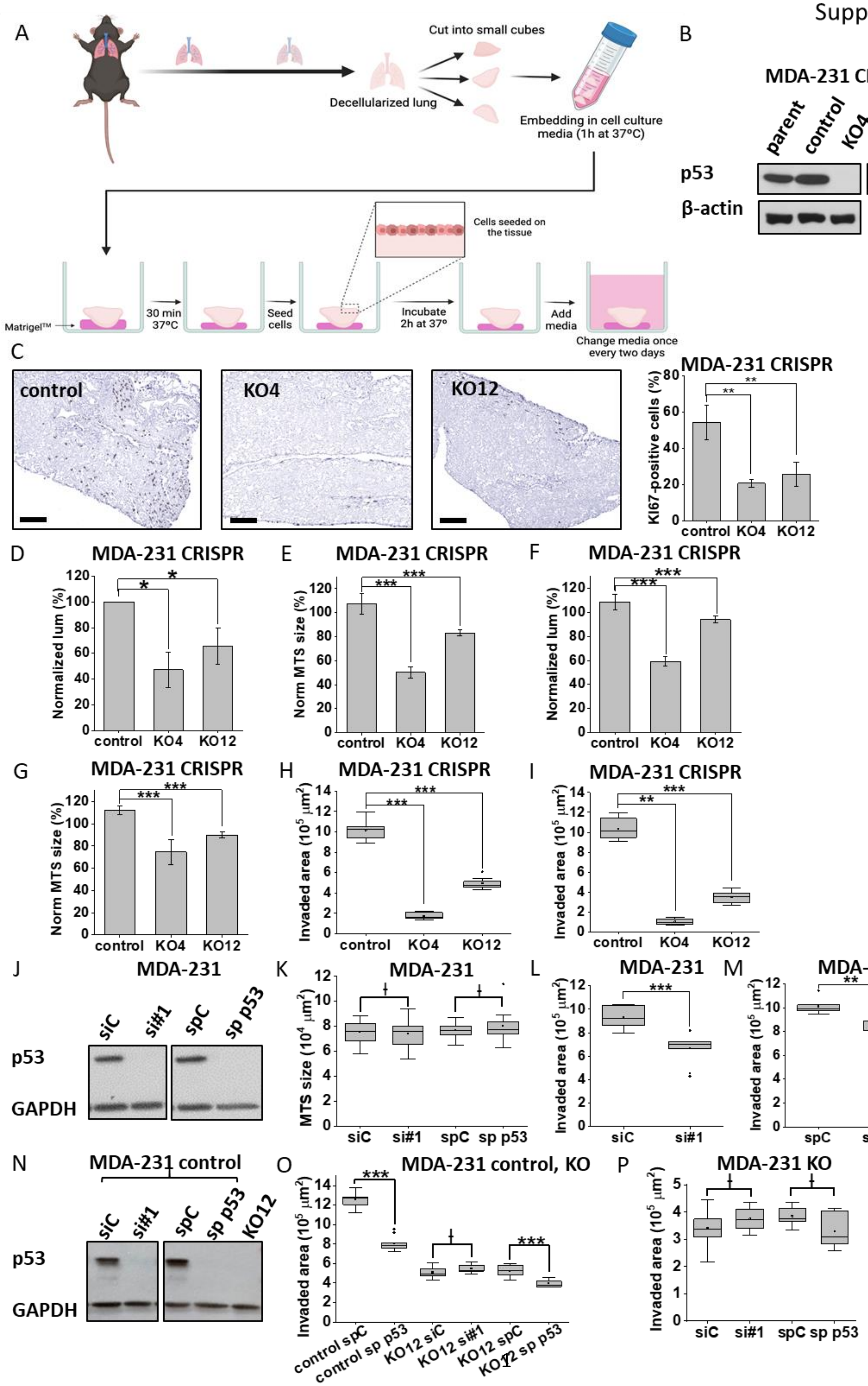

A
Cut into small cubes
Decellularized lung
Embedding in cell culture media (1h at 37°C)
Cells seeded on the tissue
Matrigel™
30 min 37°C
Seed cells
Incubate 2h at 37°
Add media
Change media once every two days
B
MDA-231 CRISPR
parent
control
KO4
KO12
p53
β-actin
C
control
KO4
KO12
MDA-231 CRISPR
KI67-positive cells (%)
D
MDA-231 CRISPR
Normalized lum (%)
E
MDA-231 CRISPR
Norm MTS size (%)
F
MDA-231 CRISPR
Normalized lum (%)
G
MDA-231 CRISPR
Norm MTS size (%)
H
MDA-231 CRISPR
Invaded area (10^5 µm^2)
I
MDA-231 CRISPR
Invaded area (10^5 µm^2)
J
MDA-231
siC
si#1
spC
sp p53
p53
GAPDH
K
MDA-231
MTS size (10^4 µm^2)
L
MDA-231
Invaded area (10^5 µm^2)
M
MDA-231
Invaded area (10^5 µm^2)
N
MDA-231 control
siC
si#1
spC
sp p53
KO12
p53
GAPDH
O
MDA-231 control, KO
Invaded area (10^5 µm^2)
control spC
control sp p53
KO12 siC
KO12 si#1
KO12 spC
KO12 sp p53
P
MDA-231 KO
Invaded area (10^5 µm^2)
siC
si#1
spC
sp p53

**Figure S1. Analysis of 3D proliferation and *ex vivo* and *in vitro* invasion of MDA-MB-231 cells with and without mutant p53.**

A) Schematics of the experimental setup for a decellularized lung tissue *ex vivo* colonization assay.

B) Western blot analysis of p53 expression in MDA-MB-231 CRISPR control and p53 KO cell lines (KO4 ad KO12) used in this study.

C) Representative images of KI67 immunohistochemistry of decellularized lung tissue (left) and KI67 score as quantified by the percentage of $KI67^+$ cells colonized on the decellularized lung tissue at Day 10 (right), (n = 4/group). Statistical significance was determined using Wilcoxon Rank Sum test. Scale bar = 50 µm.

D) MTSs were grown under ultra-low adhesion conditions for 7 days and a luminescence-based 3D proliferation assay comparing growth of MDA-MB-231 CRISPR control MTSs and CRISPR KO derivative MTSs was performed using the 3D CellTiter Glo assay. Bars represent mean luminescence normalized to the control group. n (independent biological replicates) = 3. Statistical significance was determined by two-sample t-test.

E, F) Comparison of (E) microscopy-based and (F) luminescence-based MTS size readout for the same set of MDA-MB-231 CRISPR cell spheroids. MTSs were grown under ultra-low adhesion conditions for 48 h. First, 10x DIC images were taken for MTS size analysis, subsequently the same spheroids were used for the luminescence-based 3D CellTiter Glo assay as described in D. n = 6. Within the control group the lowest value was set to 100% and all other values were normalized to that. Statistical significance was determined by two-sample t-test.

G) Microscopy-based analysis of MDA-MB-231 CRISPR MTSs at the end of the spheroid formation period of 48 h, before the onset of invasion. Spheroids were embedded in 3D collagen gels. 10x DIC images were taken immediately after completion of collagen gelation (60 min) before any visible onset of invasion. MTS size as reflected by spheroid cross section area is shown. Bars represent mean MTS size normalized to the control group. n (independent biological replicates) = 3. Statistical significance was determined by two-sample t-test.

H, I) Quantification of additional independent biological replicates of MDA-MB-231 CRISPR MTS invasion assay as shown in Fig. 1E. n = 11, 11 and 10 (H) and n = 7, 4, 7 (I) spheroids for CRISPR control, KO4 and KO12, respectively.

J) Western blot analysis of p53 knockdown efficiency in the MDA-MB-231 MTS invasion assay shown in Fig. 1F.

K) Size comparison of MDA-MB-231 MTSs under transient p53 depletion at the end of the spheroid formation period of 48 h, before the onset of invasion. Spheroids were prepared from parental cells transfected with non-targeting siRNA (siC), p53-targeting siRNA #605 (si#1), non-targeting smartpool control (spC) or p53-targeting smartpool siRNA (sp p53). 10x DIC images were taken before the onset of invasion. MTS size is presented in a box plot comprising samples from two biologically independent experiments.  n = 27, 28, 30, 23 for siC, si#1, spC and sp p53.

L, M) Additional independent biological replicates of MDA-MB-231 MTS invasion assay under transient p53 depletion as shown in Fig. 1F. n = 8, 9, 6 and 9 for siC, si#1, spC and sp p53 conditions, respectively.

N) Western Blot analysis of the p53 knockdown efficiency in the MDA-MB-231 CRISPR control MTS invasion assay shown in panel M).

O) Comparison of invaded area in MDA-MB-231 CRISPR control and KO cell lines under transient depletion of mutant p53. Analysis of invaded area from a representative MDA-MB-231 CRISPR spheroid invasion assay in 1 mg/ml collagen. n = 6 and 10 for CRISPR control transfected with spC and sp p53, and n = 9, 10, 9 and 10 for CRISPR KO12 transfected with siC, si#1, spC and sp p53.

P) Additional biological replicate of the experiment shown in panel M). n = 9, 8, 7 and 6 spheroids for CRISPR KO12 transfected with siC, si#1, spC and sp p53, respectively.

Suppl. Fig.2

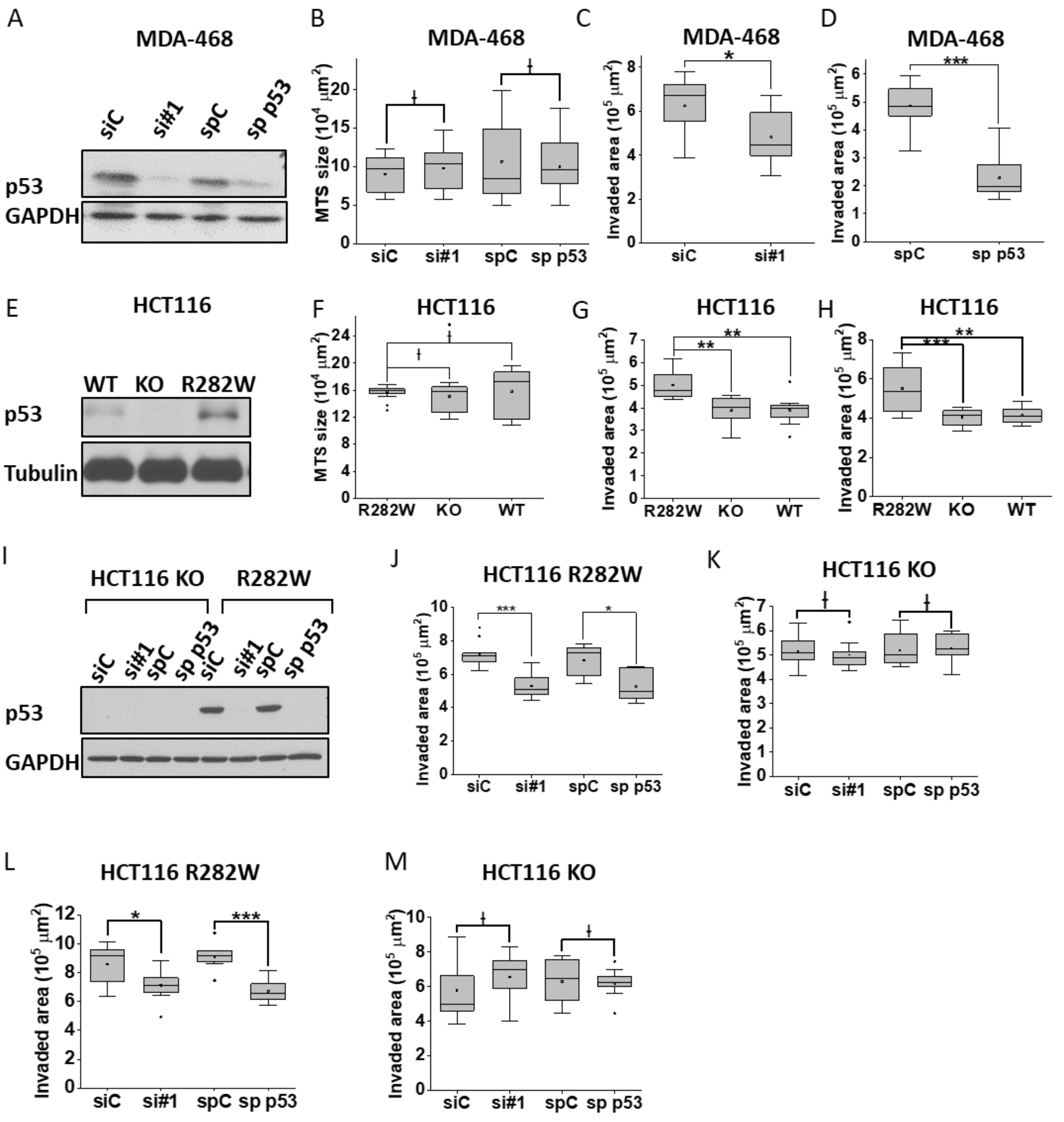
A
MDA-468
siC
si#1
spC
sp p53
p53
GAPDH
B
MDA-468
MTS size (10^4 μm^2)
siC
si#1
spC
sp p53
C
MDA-468
Invaded area (10^5 μm^2)
*
siC
si#1
D
MDA-468
Invaded area (10^5 μm^2)
***
spC
sp p53
E
HCT116
WT
KO
R282W
p53
Tubulin
F
HCT116
MTS size (10^4 μm^2)
R282W
KO
WT
G
HCT116
Invaded area (10^5 μm^2)
**
**
R282W
KO
WT
H
HCT116
Invaded area (10^5 μm^2)
***
**
R282W
KO
WT
I
HCT116 KO
R282W
siC
si#1
spC
sp p53
siC
si#1
spC
sp p53
p53
GAPDH
J
HCT116 R282W
Invaded area (10^5 μm^2)
***
*
siC
si#1
spC
sp p53
K
HCT116 KO
Invaded area (10^5 μm^2)
siC
si#1
spC
sp p53
L
HCT116 R282W
Invaded area (10^5 μm^2)
*
***
siC
si#1
spC
sp p53
M
HCT116 KO
Invaded area (10^5 μm^2)
siC
si#1
spC
sp p53

**Figure S2. 3D proliferation and invasion analysis of MDA-MB-468 and HCT116 cell lines.**

A) Western blot analysis of p53 knockdown efficiency in the MDA-MB-468 MTS invasion assay shown in Fig. 1G.

B) MTS size comparison of MDA-MB-468 under transient p53 depletion. Spheroids were prepared from cells transfected with siC, si#1, spC or sp p53 and cultured for 48 h. Box plot measuring size of spheroids as determined by area of MTS cross sections comprises samples from three biologically independent experiments. n = 35, 34, 32 and 27 for siC, si#1, spC and sp p53.

C, D) Additional independent biological replicates of MDA-MB-468 MTS invasion assay under transient p53 depletion as shown in Fig. 1G. n = 11 and 9 for siC and si#1 (C), n = 10 and 10 for spC and sp p53 (D).

E) Western Blot analysis of p53 expression in HCT116 stablecell lines used in this study.

F) Size comparison of HCT116 R282W, KO and WT spheroids at the end of the spheroid formation period of 48 h. Spheroid sizes from three biologically independent experiments are shown in box blots. n = 30, 25 and 29 for KO, WT and R282W cell lines, respectively.

G, H) Additional independent biological replicates of HCT116 MTS invasion assay as shown in Fig. 1i. n = 6, 10, 8 (G) and n = 12, 11, 8 (H) for R282W, KO and WT respectively.

I) Western Blot analysis of p53 knockdown efficiency in the HCT116 R282W MTS invasion assay shown in Fig. S1J, K.

J-M) Comparison of invaded area in HCT116 KO (K, M) and R282W (J, L) cell lines under transient depletion of mutant p53. Experiment #1 (J, K): n = 9, 8, 11 and 7 for HCT116

R282W transfected with siC, si#1, spC and sp p53 and n = 11, 10, 11 and 9 for HCT116 KO transfected with siC, si#1, spC and sp p53. Experiment #2 (L, M): n = 9, 8, 11 and 7 for HCT116 R282W transfected with siC, si#1, spC and sp p53, and n = 9, 11, 10 and 9 for HCT116 KO transfected with siC, si#1, spC and sp p53, and respectively.

Suppl. Fig.3

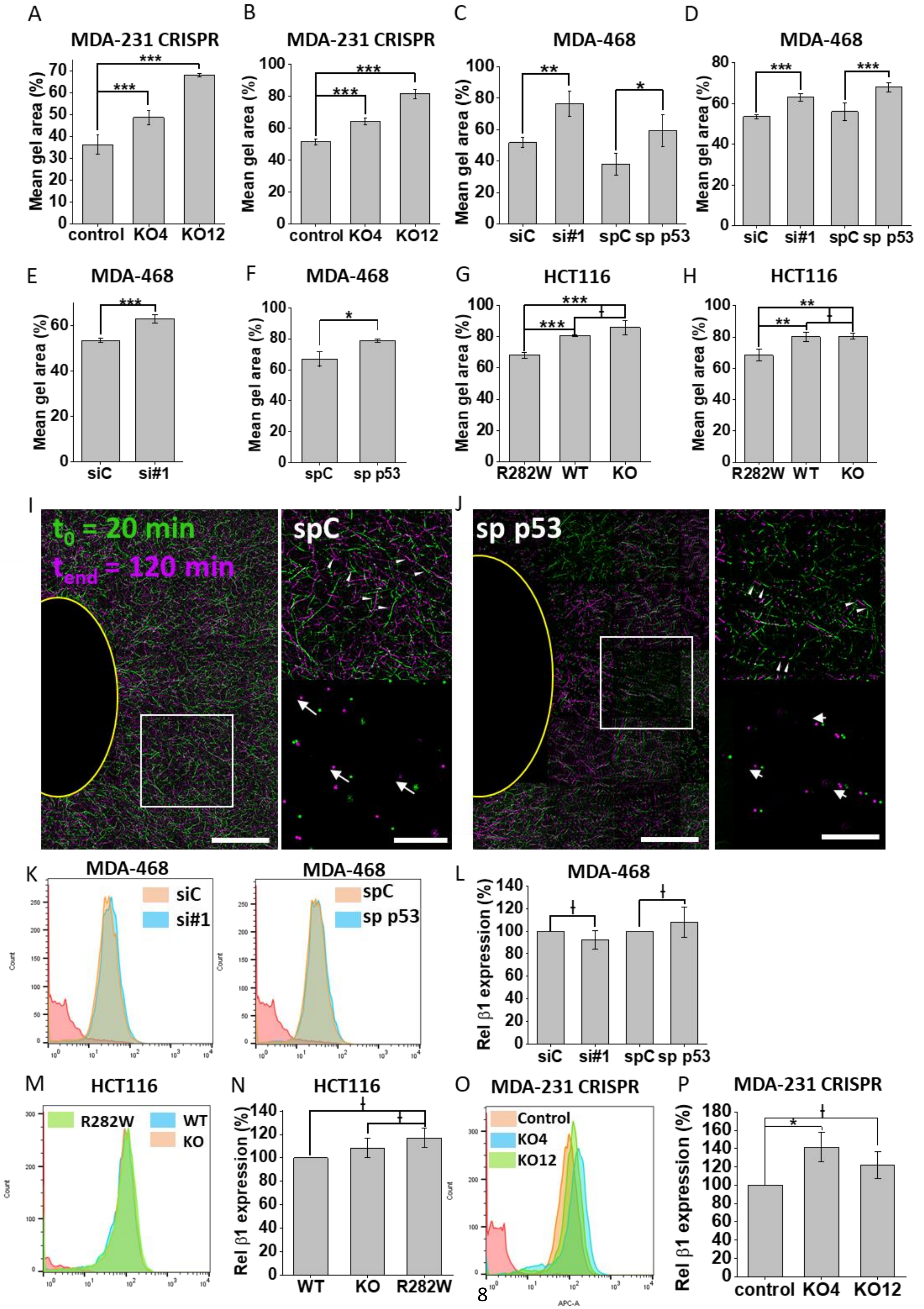
A
MDA-231 CRISPR
Mean gel area (%)
control KO4 KO12
B
MDA-231 CRISPR
C
MDA-468
siC si#1 spC sp p53
D
MDA-468
E
MDA-468
siC si#1
F
MDA-468
spC sp p53
G
HCT116
R282W WT KO
H
HCT116
I
$t_0$ = 20 min
$t_{end}$ = 120 min
spC
J
sp p53
K
MDA-468
siC
si#1
spC
sp p53
L
MDA-468
Rel β1 expression (%)
M
HCT116
R282W
WT
KO
N
HCT116
WT KO R282W
O
MDA-231 CRISPR
Control
KO4
KO12
APC-A
P
MDA-231 CRISPR
control KO4 KO12

**Figure S3. Collagen gel contraction and cell surface expression of integrin β1 receptor in cell lines with varying p53 status.**

A, B) Additional independent biological replicates of collagen gel contraction assays of MDA-MB-231 CRISPR control, KO4 and KO12 cell lines performed as described in Fig. 3B. This and following collagen contraction assays are shown as mean % gel area at the end point relative to initial gel area ± SD with statistical significance determined using two-sample unequal variance t-test. n (number of collagen matrices per condition) = 4.

C-D) Additional representative contraction assays with MDA-MB-468 cells under transient p53 depletion such as the one shown in Fig. 3C. n (number of collagen matrices per condition) = 3.

E, F) Short term collagen contraction at t = 4 h with MDA-MB-468 cells under transient p53 depletion. Two independent biological replicates are shown. n (number of collagen matrices per condition) = 6 for E) and 3 for F).

G, H) Additional collagen gel contraction assays of HCT116 KO, WT and R282W cell lines. n (number of collagen matrices per condition) = 4.

I, J) Exemplary confocal fluorescence images of cell-mediated collagen fiber re-organization at the edge of an individual MTS prepared from MDA-MB-468 cells transiently transfected with either a non-targeting (O) or p53-targeting (P) siRNA pool. MTSs were embedded into fluorescently labelled 1 mg/ml collagen I matrices and allowed to contract collagen for 2 h at 37 °C. The black void outlined in yellow indicates the edge of the MTS. Collagen imaged at the start of the experiment ($t_0$ = 20 min) is displayed in green, while collagen at the end ($t_{END}$ = 120 min) is displayed in magenta (scale bar = 50 µm). Areas defined by the white squares are shown to the right (scale bar = 20 µm) and demonstrate collagen fiber displacements (features of interest indicated by arrowheads) and displacements of embedded fluorescent beads (arrows indicate displacement direction).

K, L) Representative examples (K) and quantification (L) of FACS-based analysis of cell surface β1 integrin expression in MDA-MB-468 cells under transient p53 depletion. Histogram (K) shows distribution of fluorescence intensity of antibody-labeled β1 integrin receptors. Bar graphs (L) show average β1 integrin cell surface expression normalized to the respective control group. Data are from 3 independent biological replicates. Significance in this and following FACS analyses was determined using two sample t-test.

M, N) Representative example (M) and quantification (N) of FACS-based analysis of cell surface β1 integrin expression in stable HCT116 KO, WT or R282W cell lines. Data from 3 independent biological replicates is presented as described for K, L).

O, P) Representative example (O) and quantification (P) of FACS-based analysis of cell surface β1 integrin expression in stable MDA-MB-231 CRISPR control, KO4 and KO12 cell lines. Data from 3 independent biological replicates is presented as described above.

Suppl. Fig.4

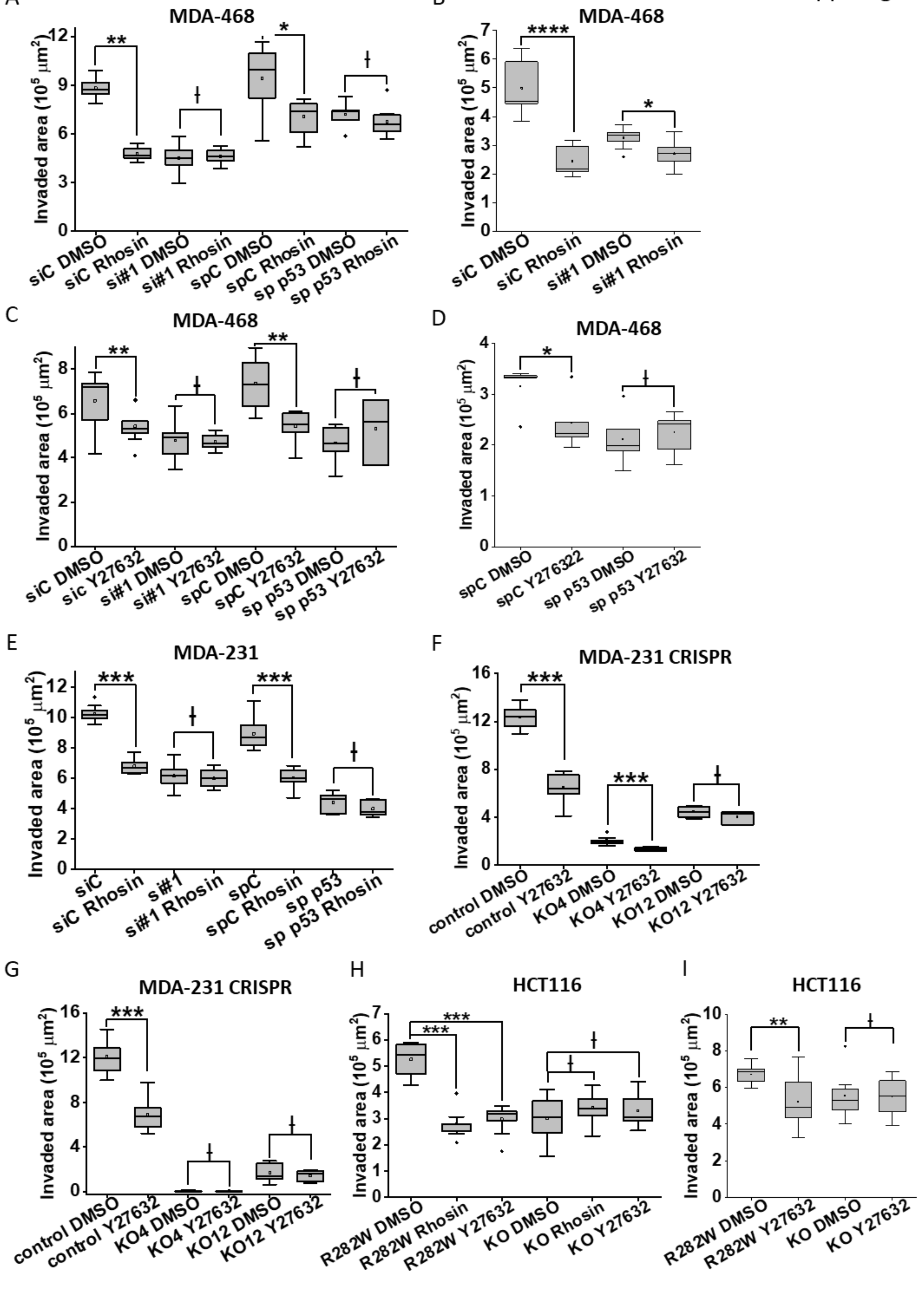
A
MDA-468
Invaded area (10^5 μm^2)
siC DMSO
siC Rhosin
si#1 DMSO
si#1 Rhosin
spC DMSO
spC Rhosin
sp p53 DMSO
sp p53 Rhosin
B
MDA-468
Invaded area (10^5 μm^2)
siC DMSO
siC Rhosin
si#1 DMSO
si#1 Rhosin
C
MDA-468
Invaded area (10^5 μm^2)
siC DMSO
siC Y27632
si#1 DMSO
si#1 Y27632
spC DMSO
spC Y27632
sp p53 DMSO
sp p53 Y27632
D
MDA-468
Invaded area (10^5 μm^2)
spC DMSO
spC Y276322
sp p53 DMSO
sp p53 Y27632
E
MDA-231
Invaded area (10^5 μm^2)
siC
siC Rhosin
si#1
si#1 Rhosin
spC
spC Rhosin
sp p53
sp p53 Rhosin
F
MDA-231 CRISPR
Invaded area (10^5 μm^2)
control DMSO
control Y27632
KO4 DMSO
KO4 Y27632
KO12 DMSO
KO12 Y27632
G
MDA-231 CRISPR
Invaded area (10^5 μm^2)
control DMSO
control Y27632
KO4 DMSO
KO4 Y27632
KO12 DMSO
KO12 Y27632
H
HCT116
Invaded area (10^5 μm^2)
R282W DMSO
R282W Rhosin
R282W Y27632
KO DMSO
KO Rhosin
KO Y27632
I
HCT116
Invaded area (10^5 μm^2)
R282W DMSO
R282W Y27632
KO DMSO
KO Y27632

**Figure S4. MTS invasion under pharmacological Rho and ROCK inhibition.**

A, B) Additional representative MDA-MB-468 spheroid invasion assays under transient p53 depletion +/- 15 µm Rhosin in 1 mg/ml collagen as shown in Fig. 3H. n = 5, 8, 8, 7, 8, 7, 7 and 8 for siC DMSO, siC Rhosin, si#1 DMSO, si#1 Rhosin, spC DMSO, spC Rhosin, sp p53 DMSO and sp p53 Rhosin in panel A; n = 7, 8, 10 and 10 for siC DMSO, siC Rhosin, si#1 DMSO, si#1 Rhosin in panel B.

C, D) MDA-MB-468 invasion assays under transient p53 depletion and pharmacological ROCK inhibition (10 µm Y27632) in low concentration collagen. n = 9, 10, 9, 10, 7, 8, 8 and 3 siC DMSO, siC Y27632, si#1 DMSO, si#1 Y27632 spC DMSO, spC Y27632, sp p53 DMSO and sp p53 Y27632 for panel C; n = 5, 5, 6, 6 for spC DMSO, spC Y27632, sp p53 DMSO and sp p53 Y27632 for panel D.

E) Additional biological replicate of MDA-MB-231 spheroid invasion assay under transient p53 depletion +/- 15 µm Rhosin in 1 mg/ml collagen as shown in Fig. 3I. n = 10, 6, 9, 7, 9, 9, 7 and 7 for siC DMSO, siC Rhosin, si#1 DMSO, si#1 Rhosin, spC DMSO, spC Rhosin, sp p53 DMSO and sp p53 Rhosin, respectively.

F, G) Quantification of two independent representative MDA-MB-231 CRISPR spheroid invasion assay under pharmacological ROCK inhibition (10 µm Y27632) in low concentration collagen. n = 9, 10, 9, 6, 4 and 3 for control DMSO, control Y27632, KO4 DMSO, KO4 Y27632, KO12 DMSO and KO12 Y27632 for panel F; n = 11, 8, 10, 8, 8 and 8 for control DMSO, control Y27632, KO4 DMSO, KO4 Y27632, KO12 DMSO and KO12 Y27632 for panel G, respectively.

H, I) Additional HCT116 spheroid invasion assays under pharmacological Rho/ROCK inhibition (15 µm Rhosin or 10 µm Y27632) in low concentration collagen. n = 8, 9, 10,

10, 7 and 10 for R282W DMSO, R282W Rhosin, R282W Y27632, KO DMSO, KO Rhosin and KO Y27632 in panel H; n = 9, 11, 8 and 8 for R282W DMSO, R282W Y27632, KO DMSO and KO Y27632 in panel I, respectively.

Supp. Fig.5

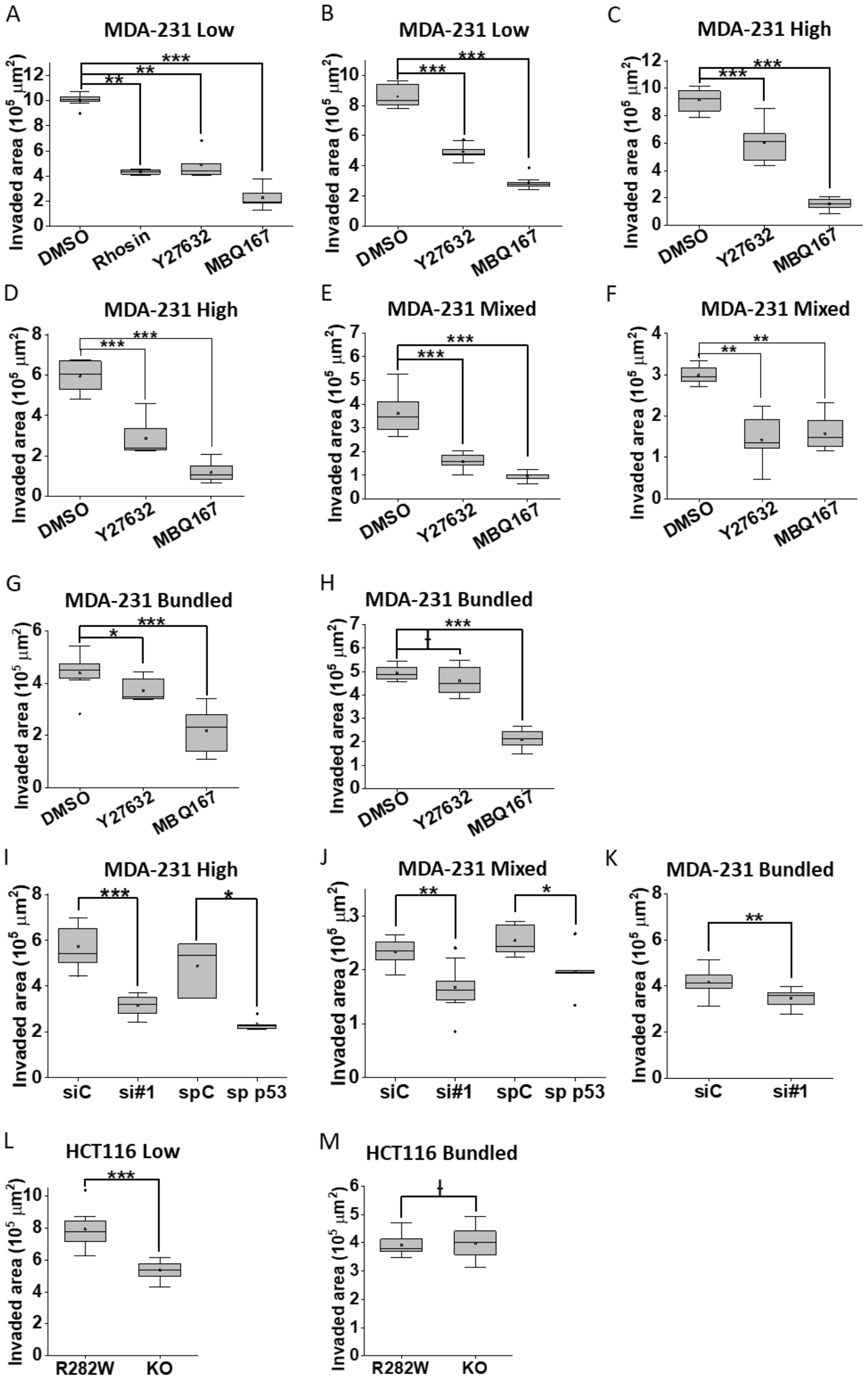
A
MDA-231 Low
Invaded area (10^5 μm^2)
DMSO
Rhosin
Y27632
MBQ167
**
**
***
B
MDA-231 Low
DMSO
Y27632
MBQ167
***
***
C
MDA-231 High
DMSO
Y27632
MBQ167
***
***
D
MDA-231 High
DMSO
Y27632
MBQ167
***
***
E
MDA-231 Mixed
DMSO
Y27632
MBQ167
***
***
F
MDA-231 Mixed
DMSO
Y27632
MBQ167
**
**
G
MDA-231 Bundled
DMSO
Y27632
MBQ167
*
***
H
MDA-231 Bundled
DMSO
Y27632
MBQ167
***
I
MDA-231 High
siC
si#1
spC
sp p53
***
*
J
MDA-231 Mixed
siC
si#1
spC
sp p53
**
*
K
MDA-231 Bundled
siC
si#1
**
L
HCT116 Low
R282W
KO
***
M
HCT116 Bundled
R282W
KO

**Figure S5. Contribution of Rho/Rac/Cdc42 pathways to MTS invasion in collagen matrices with different biomechanical properties.**

Panels A-H are biological replicates of spheroid invasion assays in biomechanically different collagen matrices. Experiments in panels A-H were performed with MDA-MB-231 cells and in panels L and M were with HCT116 R282W or KO cells.

A, B) MTS invasion under pharmacological inhibition of Rho/ROCK (Y27632) or Rac/Cdc42 (MBQ167) pathways. MTSs were formed from untreated parental cells and embedded into collagen gels supplemented with 15 µM Rhosin, 10 µM Y27632, 750 nM MBQ167 or DMSO, respectively, for the entire duration of the invasion assay (24 h). n (panel A) = 8, 4, 5 and 7 for the conditions DMSO, Rhosin, Y27632 and MBQ167; n (panel B) = 7, 9 and 10 for the conditions DMSO, Y27632 and MBQ167, respectively.

C-H) Representative MTS invasion assays in high concentration (C, D), mixed (E, F) and bundled collagen matrices (G, H) under pharmacological inhibition of ROCK (Y27632) or Rac/Cdc42 (MBQ167) pathways. n (panel C) = 7, 9, 8 for the conditions DMSO, Y27632 and MBQ167; n (panel D) = 7, 8, 7 for the conditions DMSO, Y27632 and MBQ167; n (panel E) = 8, 9, 7 for the conditions DMSO, Y27632 and MBQ167; n (panel F) = 6, 6, 7 for the conditions DMSO, Y27632 and MBQ167; n (panel G) = 9, 7, 11 for the conditions DMSO, Y27632 and MBQ167; n (panel H) = 7, 6, 7 for the conditions DMSO, Y27632 and MBQ167, respectively.

I-K) Invasion quantification of a representative MDA-MB-231 spheroid invasion assay under transient p53 depletion in I) high, J) mixed and K) bundled collagen matrices. MTSs used in both invasion assays originate from the same biological replicate. n (number of spheroids per condition) = 9, 8, 3 and 5 for siC, si#1, spC and sp p53 in high, n = 6, 9, 5 and 6 for siC, si#1, spC and sp p53 in mixed, and n = 7 and 8 for spC and sp p53 in bundled collagen matrix.

L, M) Invasion quantification of a representative HCT116 R282W and KO spheroid invasion assays in L) low concentration vs M) bundled collagen matrices.  MTSs used in both invasion assays originate from the same biological replicate. n = 8 and 9 for R282W and KO in 1 mg/ml PT, n = 10 and 9 for R282W and KO in bundled collagen.

Suppl. Fig. 6

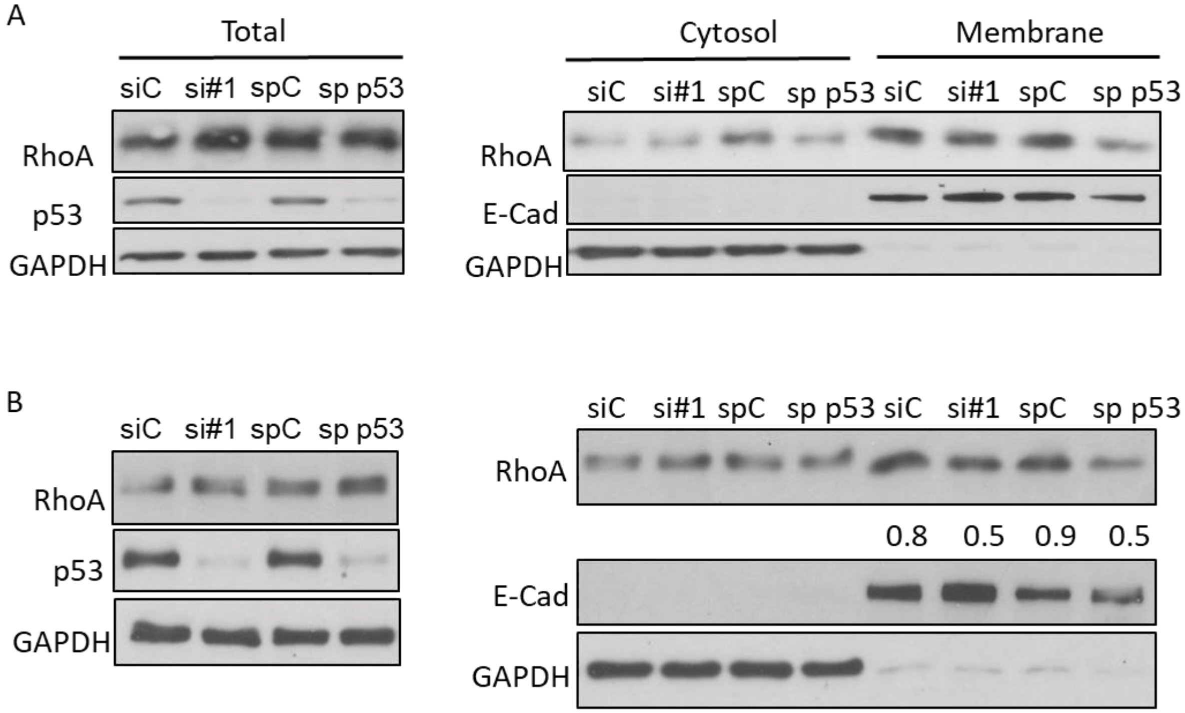
A
Total
Cytosol
Membrane
siC si#1 spC sp p53
siC si#1 spC sp p53 siC si#1 spC sp p53
RhoA
p53
GAPDH
RhoA
E-Cad
GAPDH
B
siC si#1 spC sp p53
siC si#1 spC sp p53 siC si#1 spC sp p53
RhoA
p53
GAPDH
RhoA
0.8 0.5 0.9 0.5
E-Cad
GAPDH

**Figure S6. Subcellular RhoA localization analysis by cell fractionation.**

A, B) Additional biological replicates of Western blot analysis of subcellular RhoA localization as assessed by cell fractionation in MDA-MB-468 cells under transient siRNA-mediated depletion of mutant p53 as shown in Fig. 5E. 48 h post transfection cells were seeded for 90 min on collagen-coated plates, harvested and lysates were subjected to cell fractionation. On the left are total cell lysates for the respective fractionation experiment on the right.

Suppl. Fig. 7

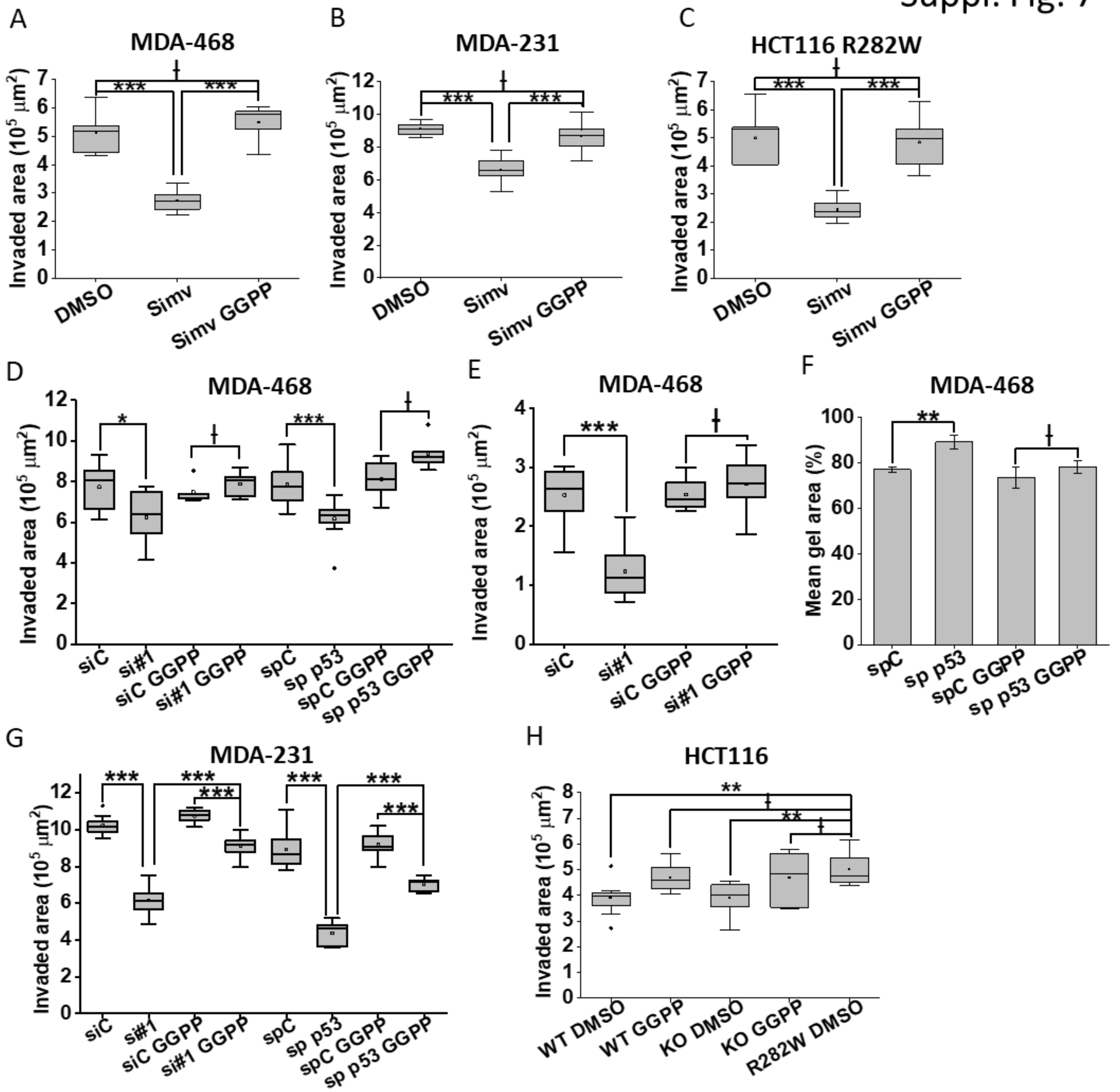
A
MDA-468
Invaded area (10^5 μm^2)
DMSO
Simv
Simv GGPP
B
MDA-231
C
HCT116 R282W
D
MDA-468
siC
si#1
siC GGPP
si#1 GGPP
spC
sp p53
spC GGPP
sp p53 GGPP
E
MDA-468
F
MDA-468
Mean gel area (%)
G
MDA-231
H
HCT116
WT DMSO
WT GGPP
KO DMSO
KO GGPP
R282W DMSO

**Figure S7. Rescue of p53-regulated MTS invasion through GGPP add-back.**

A-C) Biological replicates of MTS invasion analysis of MDA-MB-468 (A), MDA-MB-231 (B) and HCT116 (C) cells under pharmacological inhibition of the MVA pathway via simvastatin with and without exogenous supplementation of GGPP as shown in Fig. 6A-D). n (MDA-MB-468) = 10 for DMSO, Simvastatin-treated and Simvastatin/GGPP-treated conditions, n (MDA-MB-231) = 8, 11, 8 for DMSO, Simvastatin-treated and Simvastatin/GGPP-treated, n (HCT116) = 7, 9, 10 for DMSO, Simvastatin-treated and Simvastatin/GGPP-treated conditions.

D, E) MTS invasion analysis of independent biological replicates of MDA-MB-468 cells under transient depletion of mutant p53 with and without exogenous supplementation of GGPP (analogous to the assay shown in Fig. 6e, f). Fully formed MTSs prepared from cells transfected with either non-targeting (siC) or p53-targeting (si#1) siRNA were pre-treated treated with 1 mM GGPP or solvent control for 4 h in growth media and subjected to invasion assays in collagen gels supplemented with GGPP at the same concentration. n (D) = 9, 7, 7, 5, 9, 5, 10, 6 for siC untreated, sic GGPP, si#1 untreated and si#1 GGPP, spC untreated, spC GGPP, sp p53 untreated and sp p53 GGPP, respectively. n(E) = 8, 9, 8 and 7 for siC untreated, sic GGPP, si#1 untreated and si#1 GGPP, respectively.

F) Quantification of an independent biological replicate of a collagen contraction assay with MDA-MB-468 cells under transient p53 depletion with and without exogenous supplementation of GGPP (analogous to the assay shown in Fig. 6g, h). MDA-MB-468 cells were transiently transfected with either p53-targeting (sp p53) or the non-targeting control siRNAs (spC). Cells were pre-treated with 1 mM GGPP or solvent control for 4 h in growth media (2D culture) before 1 mg/ml collagen I gels +/- 1 mM GGPP loaded with

$1*10^6$ cells/ml collagen were cast. Gel area was measured before gel detachment and after 24 h of contraction. Bar graph depicts average % of contraction with standard deviation. Experiment was performed in triplicate.

G) Independent biological replicate of MTS invasion assay with MDA-MB-231 cells under transient depletion of mutant p53 with and without exogenous supplementation of GGPP (analogous to the assay shown in Fig. 6i, j). Fully formed MTSs prepared from cells transfected with either non-targeting (siC) or p53-targeting (si#1) siRNA were pre-treated treated with 1 mM GGPP or solvent control for 4 h in growth media and subjected to invasion assay in collagen gels supplemented with GGPP at the same concentration. n (d) = 10, 9, 9, 10, 9, 8, 7 and 7 for siC untreated, sic GGPP, si#1 untreated, si#1 GGPP, spC untreated, spC GGPP, sp p53 untreated and sp p53 GGPP, respectively.

H) Independent biological replicate of MTS invasion assay of HCT116 cell lines with and without exogenous supplementation of GGPP (analogous to the assay shown in Fig. 6l). Fully formed MTSs prepared from stable cell lines expressing either WT, R282W mutant or no (KO) p53 were pre-treated treated with solvent control or 1 mM GGPP for 4 h in growth media and subjected to invasion assay in collagen gels supplemented with GGPP at the same concentration. n = n = 8, 8, 10, 7 and 6 for WT DMSO, WT GGPP, KO DMSO, KO GGPP and R282W DMSO, respectively.

Suppl. Fig. 8

A

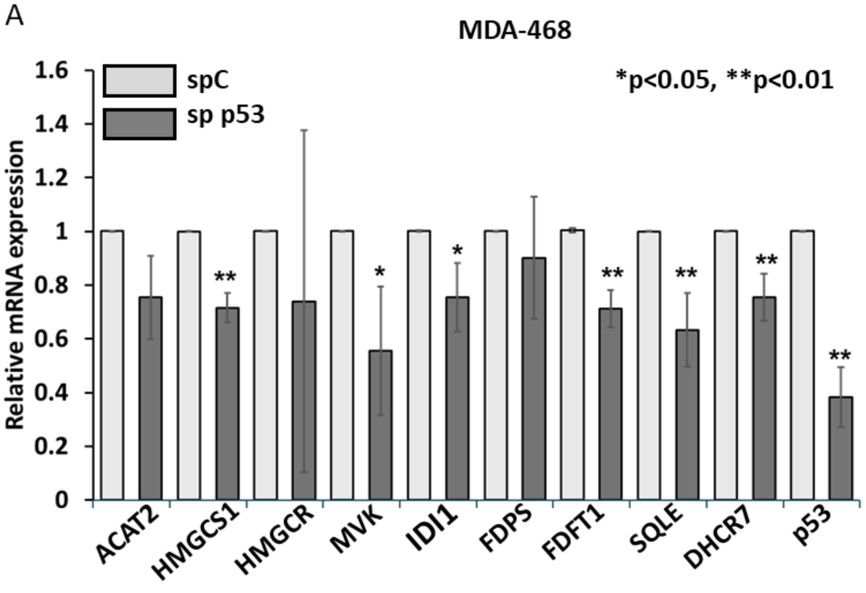


B

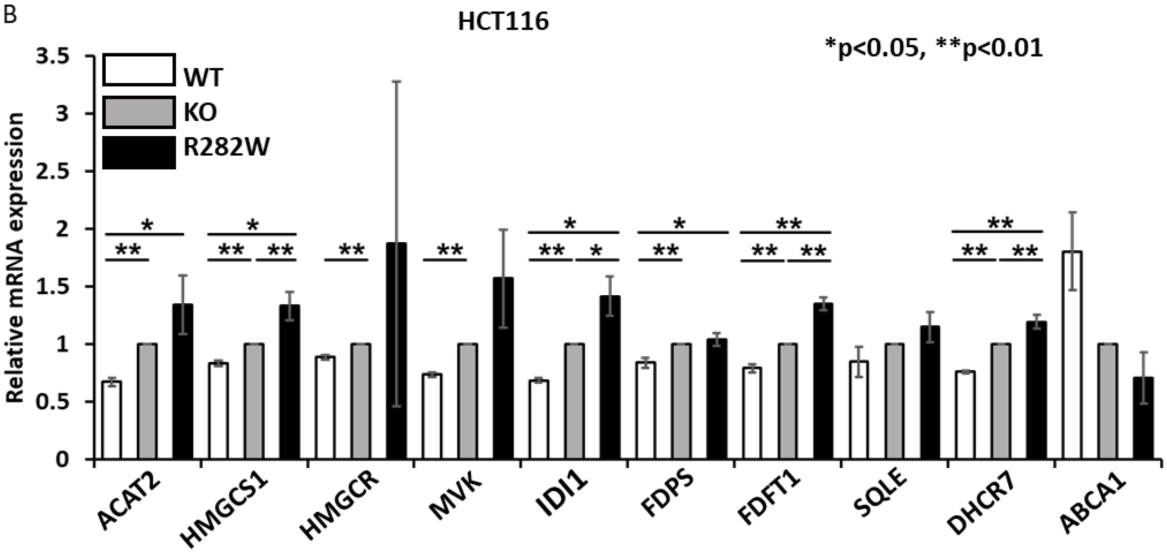

**Figure S8. Mutant p53 regulates gene expression of MVA pathway players in various cancer cell lines.**

A) RT-PCR analysis of mRNA expression of MVA pathway genes in MDA-MB-468 spheroids under transient p53 knockdown. MTSs were prepared from cells transfected with either non-targeting (spC) or p53-targeting (sp p53) siRNAs. 6 spheroids per condition were pooled for RNA extraction. Each bar represents the average from 3 biological replicates. The control was set to one and other values normalized to the control. p values were derived using two tailed t-tests from means ± SD of $n = 3$. $^*p < 0.05$, $^{**}p < 0.01$.

B) RT-PCR analysis of mRNA expression of MVA pathway genes in HCT116 WT, KO and R282W p53 spheroids. 6 spheroids per condition were pooled for RNA extraction. Each bar represents the average from 3 biological replicates. The control was set to one and other values normalized to the control. p values were derived using two tailed t-tests from means ± SD of $n = 3$ for KO and R282W, $n = 2$ for WT. $^*p < 0.05$, $^{**}p < 0.01$.

Suppl. Fig 9

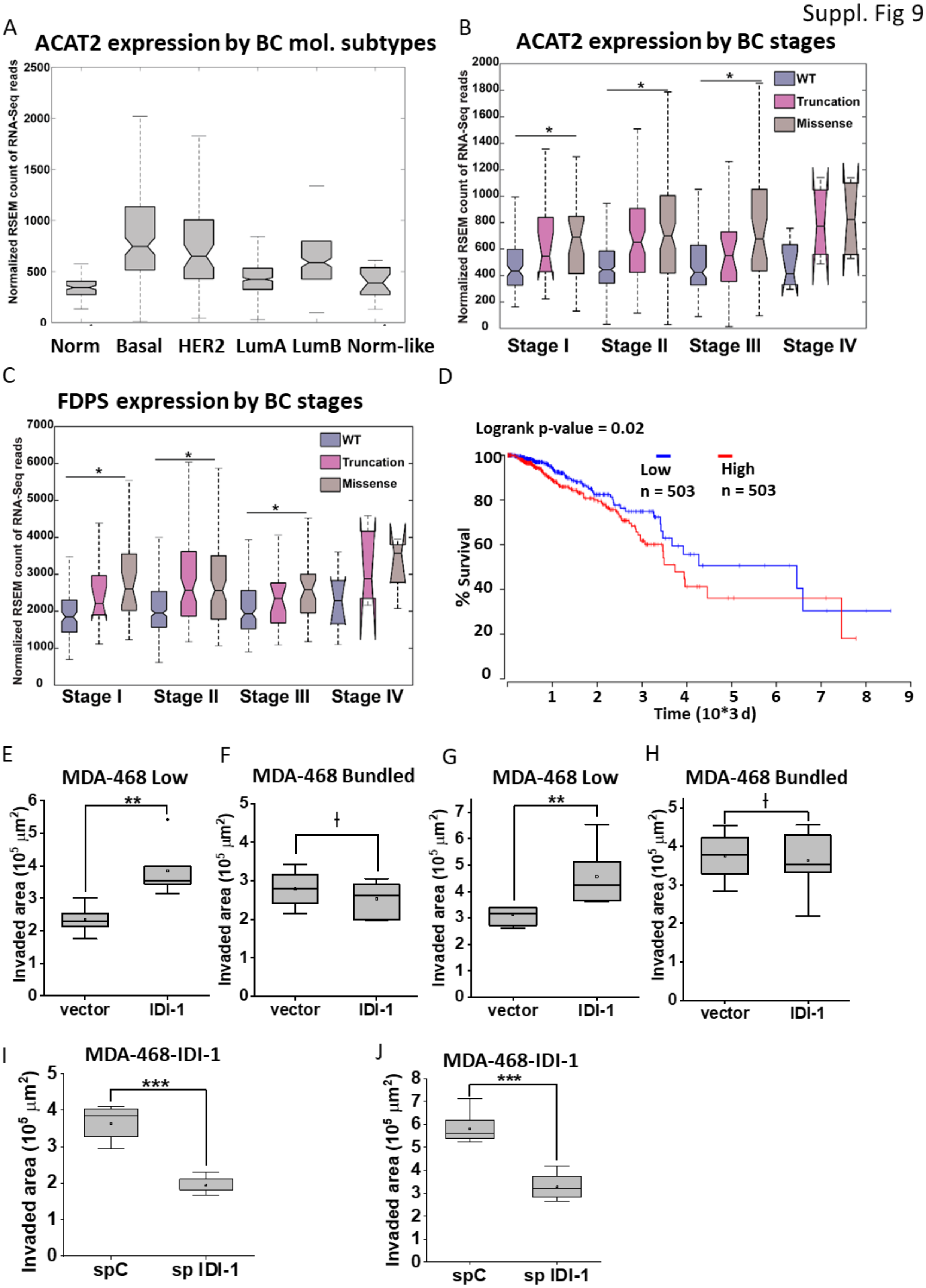
A
ACAT2 expression by BC mol. subtypes
Normalized RSEM count of RNA-Seq reads
Norm
Basal
HER2
LumA
LumB
Norm-like
B
ACAT2 expression by BC stages
WT
Truncation
Missense
Stage I
Stage II
Stage III
Stage IV
C
FDPS expression by BC stages
D
Logrank p-value = 0.02
Low
n = 503
High
n = 503
% Survival
Time (10*3 d)
E
MDA-468 Low
F
MDA-468 Bundled
G
MDA-468 Low
H
MDA-468 Bundled
Invaded area (10^5 μm^2)
vector
IDI-1
I
MDA-468-IDI-1
J
MDA-468-IDI-1
spC
sp IDI-1

**Figure S9. MVA gene expression analysis in breast cancer patients.**

A) Transcript expression of ACAT2 in TCGA breast cancer dataset (from processed RNA-seq data) stratified based on molecular subtype of the cancer. Expression of ACAT2 mRNA is significantly up regulated in tumor samples with higher frequencies of TP53 mutations compared to tumors with lower incidence of TP53 mutations and normal samples.

B, C) Transcript expression of two other genes of MVA pathway, ACAT2 (B) and FDPS (C), in TCGA breast cancer dataset (from processed RNA-seq data) stratified based the stage of the cancer and TP53 mutation status.

D) Kaplan-Meier curves based on TCGA dataset showing overall survival of cancer patients with high (red) versus low (blue) expression of IDI-1 transcript in breast cancer samples in TCGA breast cancer dataset (from processed RNA-seq data).

E-H) Additional independent repeats of the experiment shown in Fig. 7g-j. Invasion quantifications of representative MDA-MB-468-IDI-1 vs -vector control MTS invasion assays in low density vs bundled collagen matrix. The MTSs used in panels E, F and G, H originate from the same biological replicates, respectively.

I, J) Two independent biological replicates of MDA-MB-468-IDI-1 MTS invasion under transient IDI-1 depletion. Stable MDA-MB-468-IDI-1 cells were transiently transfected with either IDI-1-targeting (sp IDI-1) or non-targeting control siRNA pool (spC). Transfected cells were used for MTS formation and invasion assay. n (panel I) = 10, 9 for spC and sp IDI1; n (panel J) = 10, 13 for spC and sp IDI-1.
